# Functional screen for mediators of onco-mRNA translation specificity

**DOI:** 10.1101/2024.10.10.617637

**Authors:** Joanna R. Kovalski, Goksu Sarioglu, Vishvak Subramanyam, Grace Hernandez, Gilles Rademaker, Juan A. Oses-Prieto, Macey Slota, Nimmy Mohan, Kaylee Yiakis, Isabelle Liu, Kwun Wah Wen, Grace E. Kim, Sohit Miglani, Alma L. Burlingame, Hani Goodarzi, Rushika M. Perera, Davide Ruggero

## Abstract

Oncogenic protein dosage is tightly regulated to enable cancer cells to adapt and survive. Whether this is regulated at the level of translational control and the key factors in *cis* and *trans* remain unknown. The Myc oncogene is a central paradigm of an exquisitely regulated oncogene and a major driver of pancreatic ductal adenocarcinoma (PDAC). Using a functional genome-wide CRISPRi screen in PDAC cells, we identified activators of selective *MYC* translation through its 5’ untranslated region (5’UTR) and validated four RNA binding proteins (RBPs), including epitranscriptome modifiers. Among these RBPs, our top hit was RBM42, which is highly expressed in PDAC and predicts poor survival. Combining polysome sequencing and CLIP-seq analyses, we find that RBM42 binds and selectively regulates the translation of *MYC* and a precise, yet vital suite of pro-oncogenic transcripts, including *JUN* and *EGFR*. Mechanistically, employing IP-mass spectrometry analysis, we find that RMB42 is a novel ribosome-associated protein (RAP). Using DMS-Seq and mutagenesis analysis, we show that RBM42 directly binds and remodels the *MYC* 5’UTR RNA structure, facilitating the formation of the translation pre-initiation complex. Importantly, RBM42 is necessary for human PDAC cell growth and fitness and PDAC tumorigenesis in xenograft mouse models in a Myc-dependent manner *in vivo*. In PDAC patient samples, RBM42 expression is correlated with Myc protein levels and transcriptional activity. This work transforms our understanding of the translational code in cancer and offers a new therapeutic opening to target the expression of oncogenes.

## INTRODUCTION

The dosage of proteins encoded by onco-mRNAs is tightly regulated in order to drive cancer cell growth, adaption and survival. A key paradigm of exquisitely controlled oncogene protein expression is the master transcription factor Myc. Previous research has shown that alterations in Myc protein levels can result in a variety of phenotypic outcomes^1–3^. For oncogenic transformation, Myc levels must be maintained above homeostatic levels^4,5^. Therefore, fine tuning of Myc protein expression is critically important for initiation and maintenance of tumorigenesis. However, mechanisms of post-transcriptional regulation of Myc protein dosage, beyond control of its protein stability^6,7^, are largely unknown. A critical step in regulation of protein dosage may be transcript-specific translational control. While transcriptomic and genomic approaches have identified specific regulatory sequences within 5’UTRs, post-transcriptional control mechanisms, such as the binding of *trans*-acting RNA binding proteins have remained elusive. Similar to transcriptional control, where a defined set of proteins bind to enhancers and promoters for selective transcription, is there a similar repertoire of RNA binding proteins that act selectively in *trans* to regulate transcript-specific translational control?

In healthy cells, translation initiation of transcripts encoding for oncogenes and growth factors is tightly regulated to maintain accurate protein expression levels. However, this process is broken in cancer cells, which rely on the maintenance of high dosage of oncogenic proteins for their survival and to drive cancer development^8–12^. One possible molecular explanation may be the presence of RNA binding proteins (RBPs), which promote ribosome engagement and selectively increase protein production of oncogenic mRNAs (onco-RNAs). However, transcript-specific RBPs that may maintain onco-RNA translation at high levels to promote cancer development remain largely unknown.

Cancer cells express >1500 RBPs, which are themselves highly regulated by oncogenic signaling^13^, and therefore, the precise RBPs that selectively modulate oncogene translation have remained mostly unidentified. While efforts have been made to map the RNA interactomes of many RBPs^14^, how these RBPs interact with key RNA regulatory structures is largely unknown, due in part to a lack of *in vivo* mRNA structural analyses. Moreover, insights into how specific RBPs modulate RNA structure to regulate cellular functions are extremely limited^15^. Understanding how cancer cells co-opt RBP interaction with select *cis*-regulatory elements to precisely modulate the expression of driver oncogenes will unveil novel post-transcriptional mechanisms of gene expression and represent highly selective avenues for biomarker detection and therapeutic interventions.

Myc is constitutively expressed in >70% of human cancers and controls many of the hallmarks of cancer, such as proliferation, survival and metabolic rewiring^16^. While there has been a tremendous effort to target Myc directly and indirectly in cancer, clinically relevant Myc inhibition remains challenging^17–19^. Therefore, identifying selective regulators of *MYC* translation is a novel therapeutic opportunity. In particular, Myc is a central driver of pancreatic ductal adenocarcinoma (PDAC), which is the third-leading cause of cancer related mortality in the United States^20^. As PDAC often presents at an advanced stage with few therapies that extend 5-year survival (currently 13%)^20^, novel mechanisms to target oncogenic drivers would be of immense clinical value. High levels of Myc protein are found ∼42% of PDAC tumors, which marks the most aggressive subtype and correlates with the poorest prognosis^21–23^. Moreover, increased Myc protein expression has been shown to mediate PDAC resistance to both targeted and conventional chemotherapeutic agents^22,24–26^. Overall, unveiling the mechanisms of selective regulation of oncogene synthesis, using Myc as a paradigm, will expand the understanding of basic PDAC biology and identify new therapeutic opportunities to target this disease and other Myc dependent cancers.

Here, we develop and utilize a new functional genome-wide CRISPRi screening method to discover selective translational regulators of a key driver oncogene, Myc, in PDAC. This unbiased approach identified the regulatory network of repressors and activators of *MYC* translation genome-wide and unveiled new layers of selectivity in modulating oncogene protein expression. We identify several RBPs that specifically regulate *MYC* translation in cancer, including epitranscriptome modifiers. We focus on a top hit, a little studied RBP, RBM42, which is highly expressed in PDAC and predicts poor patient survival. We show by several genome-wide studies, including crosslinking immunoprecipitation-sequencing (CLIP-seq) and polysome sequencing, that RBM42 regulates the translation of *MYC* as well as a select, yet crucial suite of pro-oncogenic transcripts, including *EGFR* and *JUN*. Mechanistically, RBM42 binds directly to the *MYC* 5’UTR to remodel the mRNA structure, including the formation of a selective stem-loop that promotes translation. Moreover, RBM42 acts as a ribosome associated protein (RAP) and facilitates the recruitment and formation of the translation pre-initiation complex directly on the *MYC* 5’UTR, thereby increasing *MYC* mRNA translation efficiency. In patients, RBM42 expression was correlated with Myc protein levels as well as Myc transcriptional activity. *In vivo* RBM42 was necessary for PDAC tumor growth in a Myc-dependent manner. This work uncovers fundamental mechanisms and highlights new layers of specificity in translational control. Particularly, it illuminates how addiction to oncogene protein production relies upon exquisite specificity at the post-transcriptional level, generating novel cancer-specific vulnerabilities. Critically, Myc is difficult to target clinically, and therefore, selective inhibition of *MYC* translation initiation is an exciting new therapeutic opportunity for difficult to treat diseases such as PDAC.

## RESULTS

### Functional CRISPRi screen identifies selective regulators of Myc translation

The precise RBPs that selectively regulate oncogene translation have remained largely elusive, in large part due to a lack of unbiased, comprehensive methods to functionally identify candidate regulators. Illuminating the post-transcriptional mechanisms by which cancer cells precisely modulate and coordinate the specific expression of driver oncogenes can unveil new fundamental post-transcriptional mechanisms of gene expression. To identify selective regulators of *MYC* mRNA translation initiation, we used whole genome CRISPRi screening with fluorescence-activated cell sorting in human PDAC cells, where Myc plays a key role as a cancer driver (**Fig. 1a**). Specifically, we employed a functional fluorescence reporter system that enables selective monitoring of translation initiation directed by the *MYC* 5’UTR that drives the translation of a destabilized GFP, while simultaneously tracking global changes in gene expression or protein stability with a destabilized mCherry under a minimal 5’UTR. This analysis identified a few known factors involved in translation that also impinge on *MYC* translation, such as the RNA helicase eIF4A1, its co-factors eIF4H and eIF4B, as well as PCBP2, CSDE1, and the initiation site regulator eIF5A^10,27–29^, which serve as positive controls in our screen (**Extended Data Fig. 1a and Supplementary Table 1**). Importantly, we uncovered many previously unidentified putative selective activators and repressors of *MYC* translation (**Fig. 1b and Supplementary Table 1**). Functionally, Gene Ontology (GO) analysis revealed that both candidate activators and repressors of *MYC* translation were enriched for RNA binding activities, specifically RNA and mRNA binding (**Fig. 1c and Extended Data Fig. 1b**). The enrichment for DNA binding and DNA helicase factors in the potential positive regulators group may represent new transcriptional regulators that may have additional functions in interacting directly with RNA to modulate translation^30^. There is a growing body of transcription factors that have additional roles as RNA binding factors, as demonstrated by the finding that almost half of all transcription factors can bind to RNA^31^.

**Figure 1:**
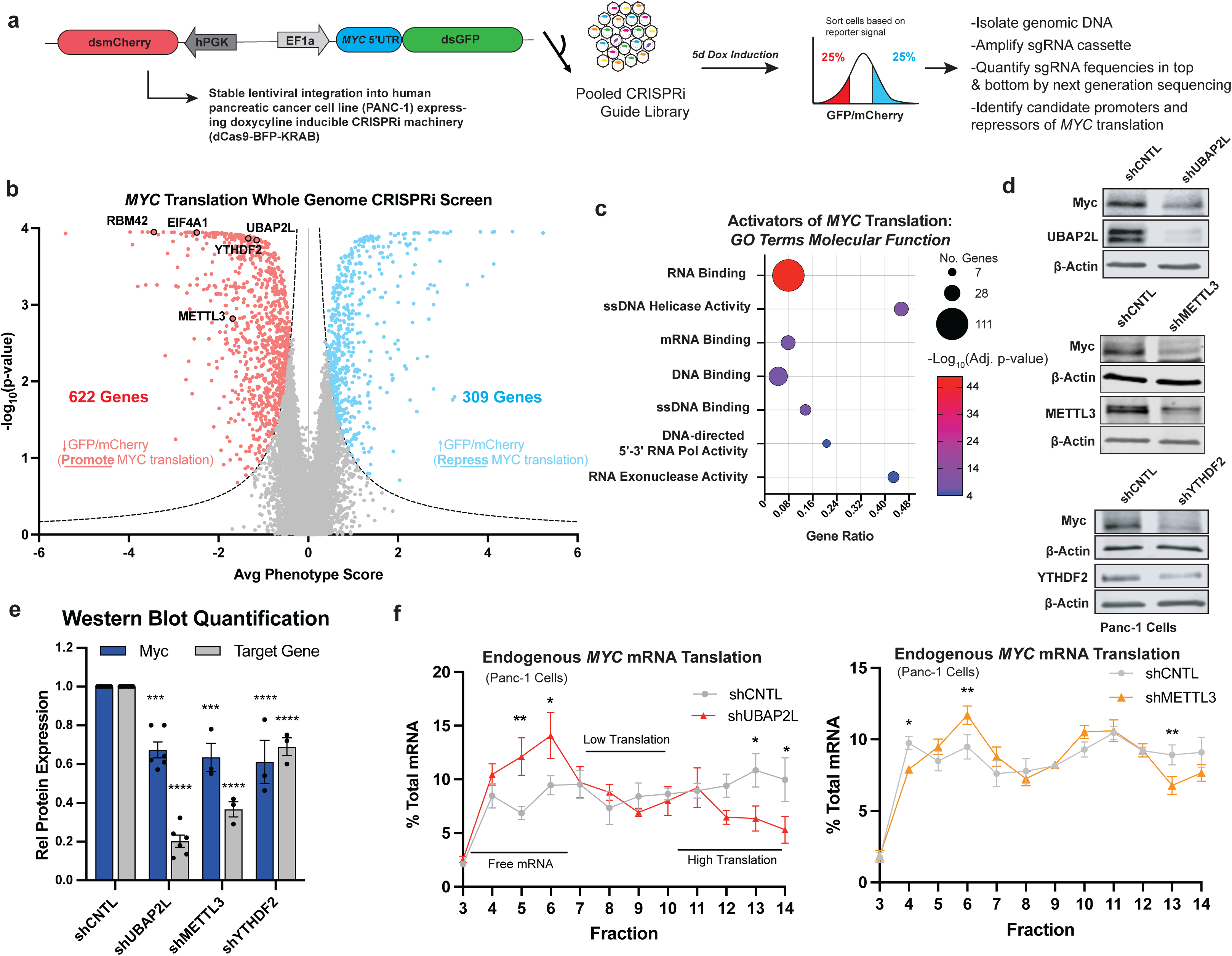
Functional CRISPRi screen identifies novel regulators of *MYC* translation. **(a)** Overview of selective *MYC* translation functional CRISPRi screen in pancreas cancer cells (Panc-1). **(b)** Volcano plot showing known and novel activators of *MYC* mRNA translation. Phenotype score is log_2_ fold change (Low GFP/mCherry sgRNA counts)/(High GFP/mCherry sgRNA counts). Significant hits have discrimination score (DS) < -1.0 (activators) or >1.0 (repressors), average phenotype score > 0.75 or < -0.75, and a p-value < 0.05. DS=average phenotype score*-log_10_[Mann-Whitney p-value]. **(c)** Gene ontology molecular function enrichment for top 300 candidate activators of *MYC* translation. Gene ratio is the fraction of genes identified within the GO term. **(d)** Western blot characterization of candidate *MYC* translational regulators’ impact on Myc protein expression in Panc-1 cells. **(e)** Quantification of candidate *MYC* translational regulators’ impact on Myc protein expression; n=3-6, mean ± SEM is shown. **(f)** qPCR analysis of *MYC* mRNA from 10-50% sucrose gradient fractionation of control or candidate translational regulator knockdown in Panc-1 cells; n=3; mean ± SEM. * p <0.05, ** p < 0.01, *** p < 0.001, **** p < 0.0001 (1- or 2-way ANOVA, uncorrected Fisher’s LSD test).

To prioritize the validation and characterization of candidate selective regulators of *MYC* translation, we focused on proteins identified to bind RNA directly in previous RNA interactome studies^32–37^. We hypothesized that these RBPs might function in *trans* at the *MYC* mRNA to directly impinge upon translation. First, we validated our CRISPRi screen results by employing single guide CRISPRi repression for select targets. This included UBAP2L, a multifunctional protein observed within stress granules^38^ whose RNA and protein levels are increased in cancer (**Extended Data Figs. 1c,d**) and was recently shown to associate with the ribosome and promote translation^39^, but with unknown selectivity and mechanism of action. Additionally, we validated m^6^A related proteins, specifically the “writer” METTL3 and “reader” YTHDF2, which emerged from our screen. m^6^A modification has been shown to regulate nearly all aspects of RNA metabolism from mRNA stability to transport and, critically, translation^40–43^. However, target specificity and the impact on translation efficiency of certain m^6^A writers and readers are highly context dependent and only beginning to be unraveled in cancer^44^. Decreased expression of UBAP2L, METTL3 and YTHDF2 in PDAC cells expressing the fluorescence reporter system demonstrated a selective decrease in the *MYC* translational reporter compared to control (**Extended Data Figs. 1e,f**). This selective change in the *MYC* translation reporter was not observed for a panel of related RBPs that were not enriched in the functional CRISPRi screen (**Extended Data Figs. 1a,g,h**).

Next, we interrogated whether these candidate regulators selectively altered endogenous Myc protein dosage. Short term depletion of UBAP2L, METTL3 or YTHDF2 in PDAC cells resulted in decreased Myc protein expression (**Fig. 1d,e**) with minor to no change in *MYC* mRNA levels (**Extended Data Fig. 1i**), implicating a translational control mechanism. To formally show that *MYC* mRNA translation was decreased with the loss of function of these hits, we employed a gold-standard method known as sucrose gradient fractionation polysome profiling. In this assay, the translation efficiency of a selected mRNA can be analyzed based on association with untranslated (free mRNA), low translation (low polysomes), or high translation (high polysomes) fractions. In these experiments, when the expression of UBAP2L, METTL3, or YTHDF2 was reduced, we observed a significant shift of the *MYC* mRNA from the polysomes (highly translating fractions) to the untranslated fractions without a decrease in total *MYC* mRNA levels (**Fig. 1f and Extended Data Figs. 1j,k**). The translation of housekeeping genes was not altered in these conditions (**Extended Data Fig. 1l**). Supporting the human clinical relevance of these *MYC* translational regulators, UBAP2L is significantly correlated with Myc protein expression levels in human PDAC quantitative proteomics (**Extended Data Fig. 1m**)^45^.

In the top 10 hits from the screen, we also identified a poorly studied RBP in the RNA Binding Motif (RBM) family, RBM42, whose role in translational control in cancer is unclear^46,47^. Intriguingly, RBM42 mRNA and protein expression are increased in PDAC compared to healthy pancreas, and high RBM42 RNA expression is correlated with decreased survival in PDAC patients^48,49^ (**Figs. 2a-c**). RBM42 is an RNA binding protein that contains a single RNA recognition motif (RRM). Previous research has predominantly focused on orthologs of RBM42 in *Toxoplasma gondii*^50^ and the fungus *Fusarium graminearum*^51^, where they play a role in pre-mRNA splicing. However, the protein sequence of the mammalian RBM42 is substantially divergent from these orthologs^50^ and its function is largely unstudied in mammalian systems. CRISPRi mediated repression of RBM42 led to a decrease in the florescent reporter driven by the *MYC* 5’UTR (**Extended Data Figs. 1e,f**). Importantly, RBM42 depletion consistently and significantly decreased endogenous Myc protein expression with minimal change in *MYC* mRNA levels across a panel of well characterized PDAC cell lines (**Fig. 2d and Extended Data Figs. 2a,b**). Furthermore, gene set enrichment analysis (GSEA) demonstrated that the “MYC targets” gene sets were the only significantly depleted Hallmark pathways with RBM42 knockdown in Panc-1 cells (adj p < 0.05; **Extended Data Fig. 2c and Supplementary Table 2**). Additionally, GSEA with transcription factor targets gene sets demonstrated significant enrichment for Myc bound genes with RBM42 depletion (**Extended Data Fig. 2c)**. RNA expression analysis via qPCR confirmed that loss of RBM42 alters the expression of direct Myc target genes (**Extended Data Fig. 2d**). These data support a predominant role for RBM42 in regulating Myc protein levels and thus modulation of Myc transcriptional target gene expression. Neither *MYC* depletion in Panc-1 cells nor enforced Myc expression in immortalized, non-transformed human pancreatic ductal epithelial cells (HPDE) significantly altered RBM42 levels, indicating that Myc does not regulate RBM42 expression (**Extended Data Figs. 2e,f**). Taken together, our functional selective translational reporter coupled with CRISPRi genetic approach identifies novel selective translational regulators of oncogenes, such as *MYC*, providing a rich regulatory network, prominently including RBPs, that may establish the multilayered complexity of translational control in cancer.

**Figure. 2:**
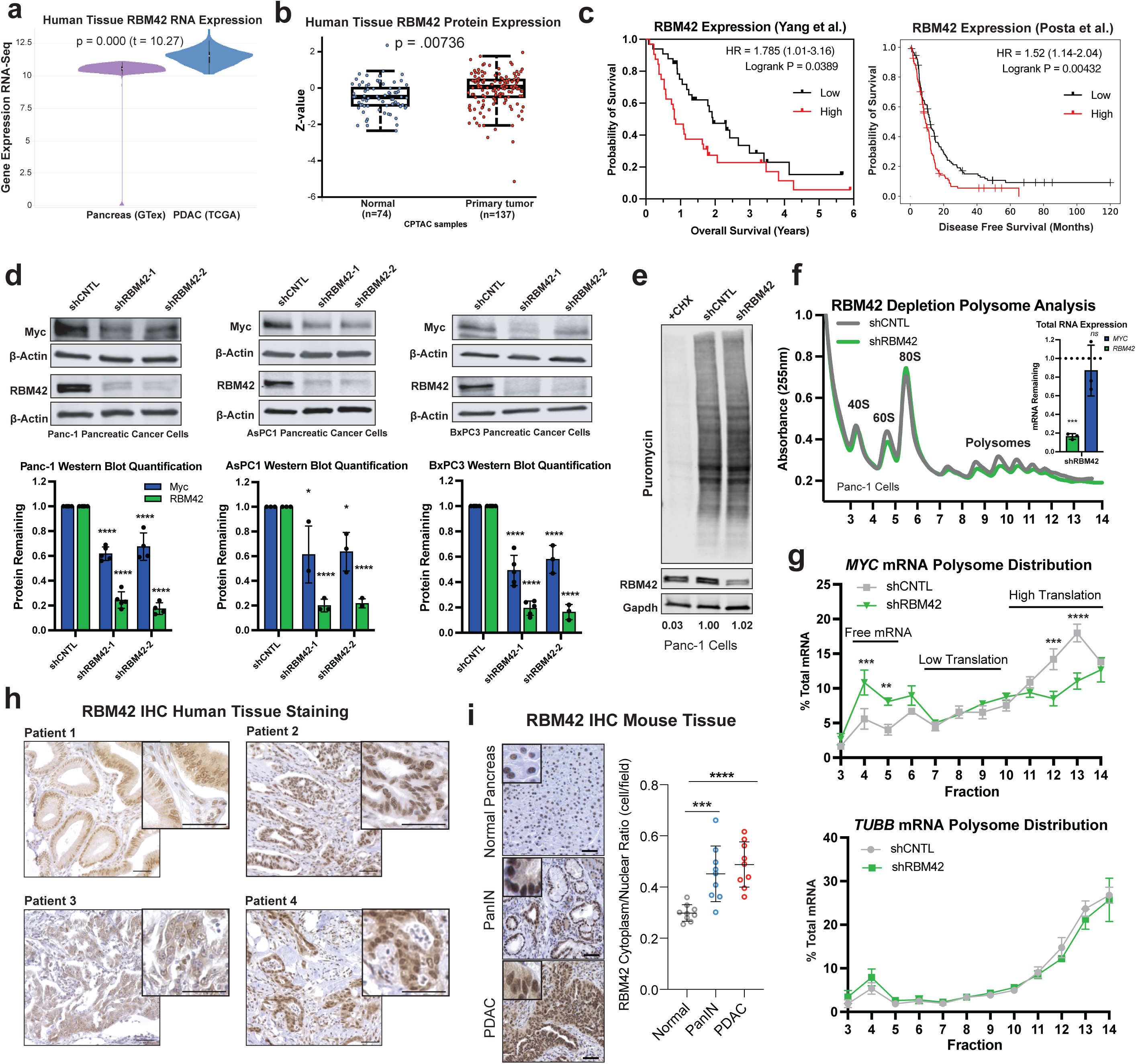
RNA binding protein RBM42 regulates *MYC* translation in pancreatic cancer. **(a)** mRNA expression of RBM42 in healthy pancreas (GTex) versus pancreatic adenocarcinoma (PDAC)^66^; Welch’s T-test. **(b)** Protein expression of RBM42 in normal pancreas versus PDAC tissues from CPTAC data set^45^; two-sided t-test. **(c)** Overall or disease-free survival correlation with RBM42 RNA expression in PDAC^48,49^. **(d)** Western blot analysis and quantification of Myc protein expression with RBM42 knockdown in a panel of pancreatic cancer cell lines. Bar graphs n=3-5; mean ± SD. **(e)** Representative western blot of puromycin incorporation assay of global translation with RBM42 depletion in Panc-1 cells. Values indicate normalized puromycin incorporation relative to shCNTL. **(f)** Representative polysome profiles for control or RBM42 knockdown in Panc-1 cells. Inset shows quantification of total RBM42 and MYC mRNA for the polysome profiling experiments in (g), mean ± SD. **(g)** qPCR analysis of *MYC* or control *TUBB* mRNA from 10-50% sucrose gradient fractionation of control or RBM42 knockdown in Panc-1 cells; n=3; mean ± SD. **(h)** RBM42 immunohistochemistry (IHC) on human PDAC tissue. Patient 1: grade 1, stage IIB; Patient 2: grade 2, stage not reported; Patient 3: grade 2, stage IV; Patient 4: grade 2, stage not reported. Scale bar = 50μm. (**i**) RBM42 IHC staining of mouse tissue and quantification of the cytoplasmic/nuclear RBM42 staining. Normal = Pft1a-Cre^+/-^, PanIN = Ptf1a-Cre^+/-^;Kras^G12D/+^, and PDAC = Ptf1a-Cre^+/-^;Kras^G12D/+^;Trp53^flox/+^; n = 3 mice/condition, 3 fields/animal; mean ± SD. Unpaired t-test. Scale bar = 50μm. * p < 0.05, ** p < 0.01, *** p <0.001, **** p < .0001, *ns* = not significant (1- or 2-way ANOVA, uncorrected Fisher’s LSD test).

### RBM42 is a new regulator of *MYC* translation

To test the sufficiency of RBM42 to regulate Myc protein levels, we expressed RBM42 in immortalized, non-transformed human pancreatic ductal epithelial cells (HPDE) commensurate with the expression observed in PDAC cells. In this setting, RBM42 alone was sufficient to increase Myc protein expression, while *MYC* mRNA expression was unchanged (**Extended Data Fig. 2g**). While the data implicate RBM42 in the selective regulation of *MYC* mRNA translation control, we next examined the impact of RBM42 on global protein synthesis. Loss of RBM42 did not alter global translation levels in puromycin incorporation assays across several PDAC lines (**Fig. 2e and Extended Data Fig. 2h**). In support of the hypothesis that RBM42 has a specific role in translation, the overall polysome profile upon RBM42 loss of function was not altered (**Fig. 2f**). However, RBM42 depletion selectively decreased the translation of the *MYC* mRNA but did not affect the translation of the control mRNA, *TUBB* (**Fig. 2g**). Myc is known to be regulated at the post-translational level^6,7^. Therefore, to exclude the impact of RBM42 on Myc protein stability, we conducted a cycloheximide time course experiment in PDAC cells. RBM42 loss did not decrease Myc protein stability (**Extended Data Fig. 3a**).

RBM42 was identified as an accessory factor in the human U4/U6.U5 tri-snRNP cryo-EM structure^52^. However, reduced RBM42 levels did not alter splicing of *MYC* or a control (*B2M*) mRNA by qPCR in Panc-1 cells (**Extended Data Fig. 3b**). As a previous study implicated RBM42 in splicing, we conducted global splicing analysis using rMATS on triplicate paired end RNA-sequencing of Panc-1 cells with control or RBM42 knockdown. Compared to the number of splicing events, including retained introns, skipped exons, mutually exclusive exons and alternative 5’ or 3’ splice site usage, observed with the loss of core, canonical spliceosome proteins (e.g. U2AF2, SRSF1, MAGOH) or snRNAs^14,53^, RBM42 depletion exhibited very minimal altered splicing events and no splicing alterations of the *MYC* mRNA (**Extended Data Fig. 3c and Supplementary Table 3**). To substantiate a new cytoplasmic translational control function for RBM42, we stained a panel of 19 human PDAC tissues, representing a broad spectrum of grades (1-3) and stages of disease (IB to IV). RBM42 immunohistochemistry (IHC) showed cytoplasmic and nuclear RBM42 localization across all patient tissues (**Fig. 2h and Extended Data Fig. 3d**). To examine RBM42 subcellular distribution over the course of oncogenic transformation, we performed RBM42 IHC on normal pancreas, pre-cancerous pancreatic intraepithelial neoplasias (PanINs) and PDAC tissue from a mouse model that recapitulates human PDAC. Strikingly, RBM42 cytoplasmic localization increased dramatically in both PanINs and PDAC tissues compared to normal pancreas, suggesting that RBM42 may regulate *MYC* translation from early stages of tumorigenesis (**Fig. 2i**). Finally, subcellular fractionation of a panel of PDAC cell lines showed that a portion of RBM42 is localized in the cytoplasm, further supporting RBM42’s role in the cytoplasm (**Extended Data Fig. 3e**). These data support a novel function for cytoplasmic localized RBM42 in promoting the selective translation of the *MYC* mRNA to maintain Myc protein levels in PDAC.

### RBM42 is necessary for PDAC cell viability through Myc

RBM42 RNA and protein expression is significantly increased in human PDAC (**Figs. 2a,b**), which suggests that high levels of RBM42 are favorable for carcinogenesis. Moreover, Myc is a potent driver of many facets of tumorigenesis. Therefore, we sought to functionally link RBM42-dependent regulation of Myc protein expression to key hallmarks of cancer. Depletion of RBM42 in several different PDAC lines resulted in dramatically decreased cell growth and cell viability, and diminished colony formation (**Figs. 3a,b and Extended Data Fig. 4**). In parallel, PDAC cells with reduced RBM42 expression were challenged with anchorage independent growth in soft agar, which mimics tumor growth *in vivo*. RBM42 loss significantly reduced the ability of PDAC cells to form colonies in three dimensions (**Fig. 3c**). To determine whether the observed reduction in cell viability is Myc dependent, we enforced expression of Myc lacking the 5’UTR. Myc expression was sufficient to rescue the reduced cell viability associated with RBM42 loss in two independent PDAC lines (**Figs. 3d,e**). Therefore, RBM42 is a *bona fide* oncogenic factor required for the maintenance of PDAC through Myc.

**Figure. 3:**
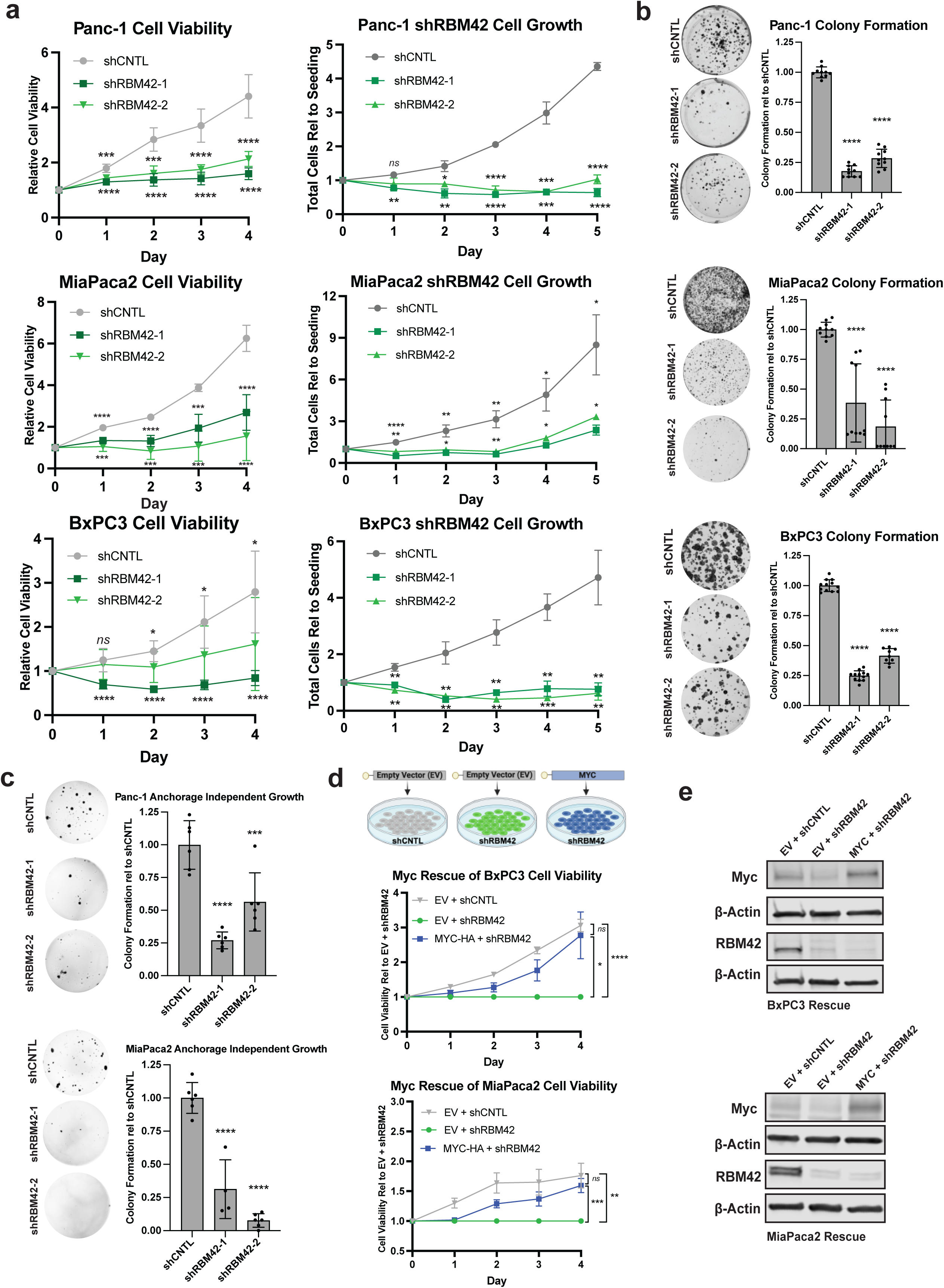
RBM42 is necessary for pancreatic cancer cell growth through Myc. **(a)** Pancreatic cancer cell line viability or cell growth time course with control (CNTL) or RBM42 shRNA knockdown using Cell Titer Glo Assay normalized to day 0 n= 6 (Panc-1 & MiPaca2) or 9 (BxPC3) or total live cell counting (trypan blue negative) relative to the number of cells seeded (n = 4-6 replicates). Unpaired t-test. **(b)** Two-dimensional colony formation with control or RBM42 depletion stained with crystal violet; n=10-12. **(c)** Anchorage independent growth in soft agar with control or shRBM42 stained with MTT to visualize live colonies; n=4-6. **(d)** Myc coding sequence (CDS) rescue of shRBM42 induced loss of PDAC cell viability. To calculate the cell viability relative to EV + shRBM42, all readings were normalized to their day 0 values to account for cell seeding. Then, for each timepoint, all conditions were compared to the average of knockdown (EV + shRBM42) viability readings for that day to calculate the fold rescue.; n=10 (BxPC3) or 6 (MiaPaca2), significance at day 4 mean ± SEM. Diagram created with BioRender.com. **(e)** Representative western blot of the Myc and RBM42 expression for cell viability rescue experiments. Graphs are mean ± SD unless noted. Ordinary 1-way ANOVA with Dunnett’s Multiple Comparison Test unless otherwise indicated. * p < 0.05, ** p < 0.01, *** p <0.001, **** p < .0001, ns = not significant.

### RBM42 Regulates an Oncogenic Translational Program

RBM42 regulates the protein dosage of Myc in PDAC cells, but it is unclear how RBM42 increases the translation efficiency of the *MYC* mRNA or if RBM42 may regulate the translation of additional oncogenic transcripts. Therefore, we assayed transcriptome-wide cytoplasmic RBM42 binding using crosslinking immunoprecipitation sequencing (CLIP-seq). We found that RBM42 directly bound the *MYC* 5’UTR with a substantial enrichment in the 5’UTR compared to the rest of the transcript (**Figs. 4a,b and Extended Data Fig. 5a**). Furthermore, CLIP-qPCR demonstrated significantly enriched binding of RBM42 to the mature *MYC* mRNA in the cytoplasmic fraction, supporting RBM42’s role in translational regulation of *MYC* (**Extended Data Figs. 5b,c**). Additionally, the CLIP analysis revealed that RBM42 significantly bound the 5’UTRs of 2312 genes, which were functionally enriched for cancer associated pathways (**Extended Data Fig. 5d and Supplementary Table 4**). The CLIP-seq data support RBM42’s direct regulation of *MYC* translation, while also implicating RBM42 in the regulation of a broader gene expression program.

**Figure. 4:**
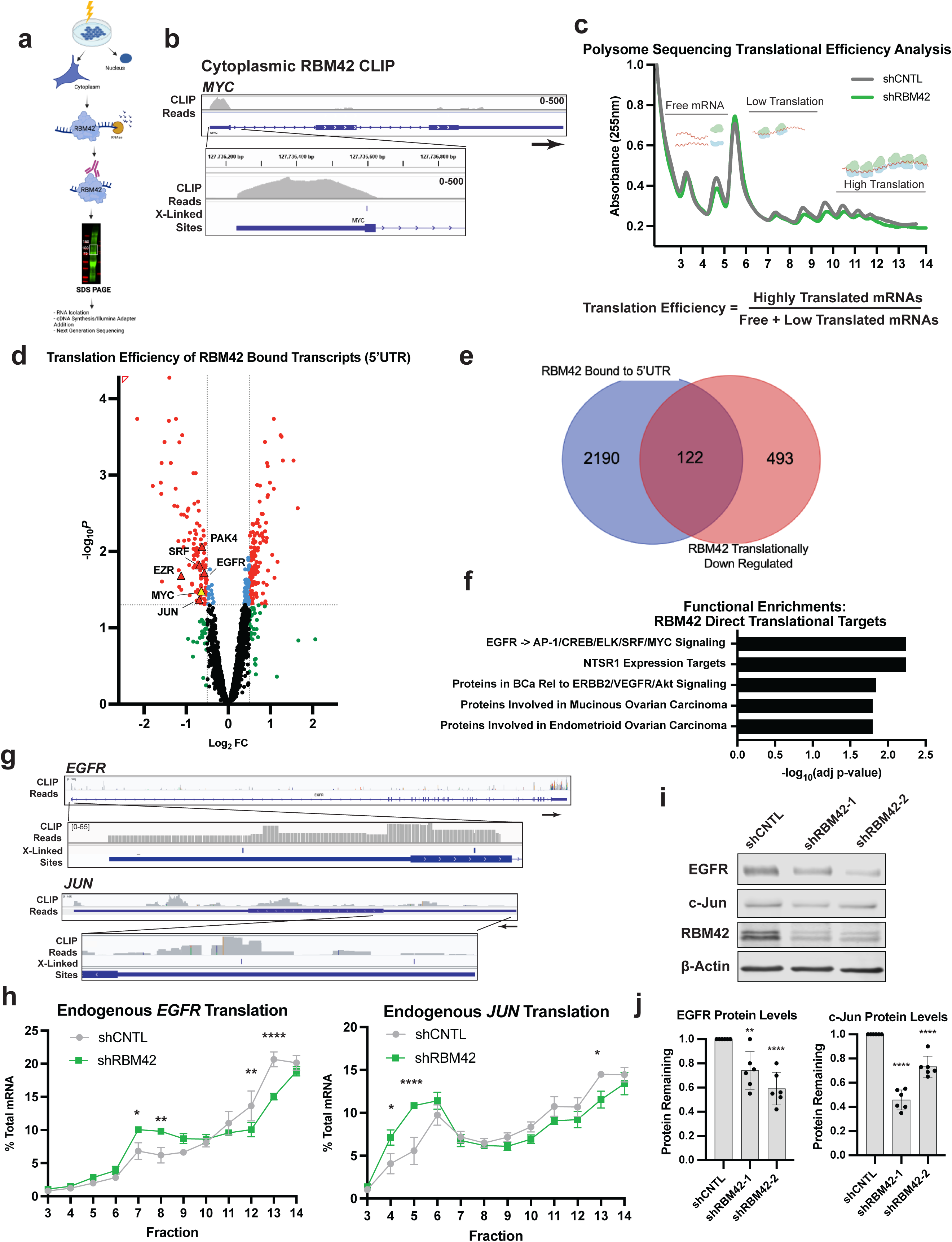
RBM42 directly binds and regulates the translation of a pro-oncogenic program. **(a)** Scheme of the endogenous cytoplasmic RBM42 CLIP-seq workflow in Panc-1 cells. Created with BioRender.com. **(b)** CLIP reads and crosslinked induced mutation sites (CIMS) on the *MYC* mRNA, inset of 5’UTR (CIMS FDR < 0.05). **(c)** Diagram of polysome sequencing and translational efficiency calculation in Panc-1 cells. **(d)** Volcano plot of translational efficiency of genes bound by RBM42 in the 5’UTR, key pro-oncogenic genes highlighted. **(e)** Venn diagram of RBM42 5’UTR bound CLIP genes (FDR < .05) and RBM42 translationally downregulated genes (adj p-value < 0.05 and FC < 0.75 [log_2_FC < -0.42]). **(f)** Functional enrichment of Elsevier Pathways of CLIP/Poly-Seq overlapping genes. **(g)** CLIP tracks for additional RBM42 bound and translationally regulated targets, *EGFR* and *JUN*. Arrow indicates the orientation of the gene, inset of 5’UTR (CIMS FDR < 0.05) **(h)** qPCR analysis of *EGFR* or *JUN* mRNA from 10-50% sucrose gradient fractionation of control or RBM42 knockdown in Panc-1 cells; n=3; mean ± SEM. (2-way ANOVA, uncorrected Fisher’s LSD test). **(i)** Representative western blots of EGFR and c-Jun protein expression with RBM42 depletion. **(j)** Protein quantification for EGFR and c-Jun with RBM42 depletion in Panc-1 cells (n = 6), unpaired t-test. All graphs mean ± SD unless noted. * p < 0.05, ** p <0.01, **** p < .0001.

This prompted further analyses to hone in on the genes that are both directly bound and translationally regulated by RBM42 that would comprise a coordinated RBM42 translational program. To this end, we employed an unbiased genome-wide translational profiling approach coupled to RNA sequencing (Polysome-sequencing) (**Fig. 4c**). Polysome-sequencing enables direct measurement of ribosome density per mRNA independent of RNA expression changes, providing a direct estimate of translation efficiency. We calculated the relative translational efficiency (TE) for each mRNA, which is the enrichment of a given transcript in the heavy polysome fractions (highly translated) compared to the sum of free/ribonucleoproteins (RNP), 80S and light polysome (untranslated and low translation) fractions (**Fig. 4c and Supplementary Table 5**). Overlapping the transcripts with RBM42 bound in their 5’UTRs with those exhibiting decreased translational efficiency upon RBM42 depletion identified 122 genes (**Figs. 4d,e**). Interestingly, these genes were enriched for many cancer signaling pathways and prominent pro-oncogenic proteins (**Figs. 4d,f**). First, we focused on the oncogenes epidermal growth factor receptor (*EGFR*) and the transcription factor c-Jun (*JUN*). RBM42 bound the 5’UTR of both *EGFR* and *JUN* (**Fig. 4g**) without affecting their splicing (**Supplementary Table 3**). Using sucrose gradient fractionation polysome profiling, we observed that RBM42 depletion shifted both the *EGFR* and *JUN* mRNA from the highly translated to the untranslated fractions, and concomitantly decreased EGFR and c-Jun protein levels (**Figs. 4h-j**). Additionally, RBM42 altered the translation efficiency and, thereby the protein expression, of other key genes involved in a variety of cancer processes, such as *PAK4*, involved in signal transduction, *EZR* implicated in cytoskeletal dynamics, and the transcription factor, *SRF* (**Extended Data Figs. 5e**-g). We further validated RBM42 regulation of translation initiation for this set of genes (*EGFR*, *JUN*, *SRF*, *PAK4* and *EZR*) using luciferase reporters whose translation are driven by each genes’ 5’UTR (**Extended Data Fig. 5h)**. Overall, we uncovered that RBM42 binds and promotes the translation of a select, yet critical suite of transcripts, illuminating a novel means by which an RNA binding protein can coordinate the expression of a specific network of pro-oncogenic factors in pancreatic cancer.

### RBM42 remodels the *MYC* 5’UTR structure to regulate translation

As RBM42 directly binds the *MYC* 5’UTR to increase translation efficiency, we next investigated the molecular mechanisms by which RBM42 controls *MYC* translation efficiency. We tested whether RBM42 may modulate specific RNA elements within the endogenous *MYC* 5’UTR RNA structure in Panc-1 PDAC cells. We performed *in vivo MYC* 5’UTR RNA structure analysis using dimethyl sulfate (DMS) treatment followed by targeted sequencing. DMS modifies unpaired adenines and cytosines, which cause mutations during reverse transcription; thus, the mutation frequency is an estimate of nucleotide accessibility and the likelihood of base pairing^54^. The endogenous *MYC* 5’UTR mRNA structure illuminated new, previously unknown aspects of this highly complex RNA structure within cancer cells. Unlike the previous *in vitro* determined structure^55^, the endogenous RNA contains three stems, one of which contains a large open loop structure at the 3’ end (referred to here as Stem III), juxtaposed to the main start codon (**Figs. 5a,b and Supplementary Table 6**). To uncover where RBM42 is most likely to bind to the *MYC* 5’UTR structure, we used crosslink-induced mutation sites (CIMS) analysis^56^ of the RBM42 CLIP-seq data, which identifies statistically significant nucleotide mutations introduced by the reverse transcriptase encountering protein-RNA crosslinks during cDNA synthesis. This analysis identified an RBM42 binding site at U369 in the large loop in the Stem III structure (**Fig. 5b**, red arrow).

**Figure. 5:**
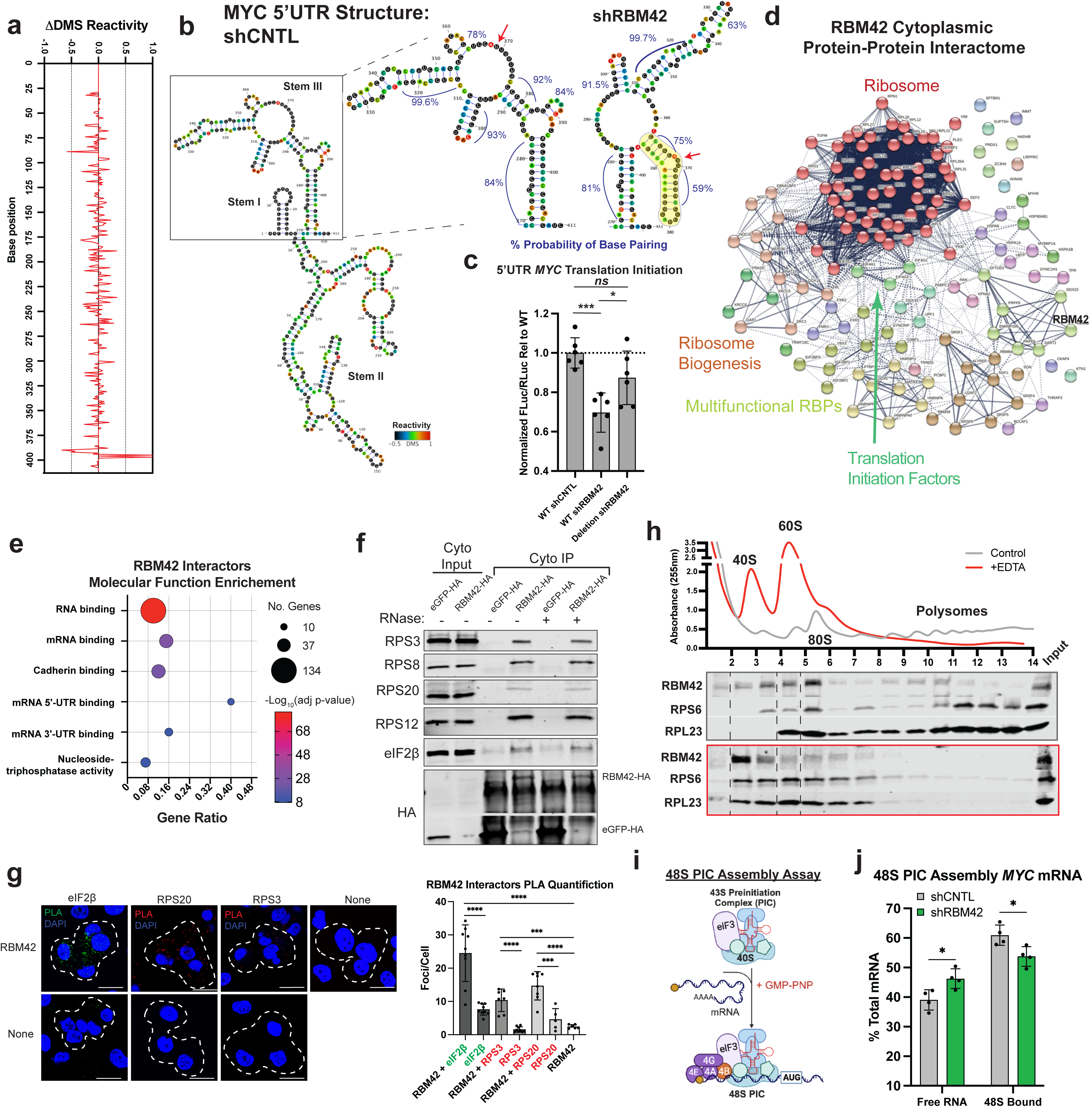
RBM42 remodels the structure of the *MYC* 5’UTR and acts as a RAP to promote efficient translation initiation. **(a)** Plot of the difference in dimethyl sulfate (DMS) reactivities of the *MYC* 5’UTR between shRBM42 versus shCNTL in Panc-1 cells. **(b)** DMS-Seq determined full length *MYC* 5’UTR RNA structure in control cells, zoom in on Stem III comparing shCNTL vs. shRBM42. Numbers indicate the average probability of the base pairing in the specified region. Red arrow indicates the predicted RBM42 bound nucleotide as show in Fig. 4b. **(c)** Dual luciferase assay with the *MYC* 5’UTR WT or with deletion of bases 363-394, highlighted in Fig. 5b, in Panc-1 cells. Luciferase values normalized to RNA expression levels, n=6. **(d)** IP followed by mass spectrometry identified endogenous RBM42 cytoplasmic protein-protein interactome in Panc-1 cells. Relevant functional groupings labeled. **(e)** Molecular function gene ontology enrichment of RBM42 interacting proteins. Gene ratio is the fraction of genes identified within the GO term. **(f)** Representative IP-western blots of eGFP-HA or RBM42-HA from the cytoplasmic fraction in the presence or absence of RNaseA treatment with 40S ribosomal subunits (RPS) and eIF2β (component of the 43S PIC). (**g**) Proximity ligation assay images and quantification of foci per cell in Panc-1 PDAC cells probing for RBM42 with either eIF2β, RPS20 or RPS3. None indicates a single primary antibody was used. Dotted lines indicate the outline of the cells, n ≥ 5 fields/condition. (**h**) Representative western blot of fractions from Panc-1 untreated or EDTA treated cytoplasmic lysate separated on a 10-50% sucrose gradient; small ribosomal subunit member, RPS6, and large ribosome subunit protein, RPL23. **(i)** Diagram showing the 48S pre-initiation complex formation assay. Created with BioRender.com. **(j)** Quantification of *MYC* mRNA in the free RNA versus 48S bound RNA fractions from a 10-30% sucrose gradient fractionation of control or shRBM42 knockdown Panc-1 lysate, n=4. All graphs mean ± SD. * p < 0.05, *** p <0.001, **** p <0.0001, ns = not significant (Unpaired t-test).

We reasoned that RBM42 binding may alter the *MYC* 5’UTR structure to a conformation that promotes increased translation initiation. Interestingly, the *MYC* 5’UTR was remodeled upon RBM42 loss with the most prominent and significant changes in Stem III, but with smaller alterations in other regions (**Fig. 5a**). Specifically, the region around the predicted RBM42 direct binding site and proximal to the AUG start codon exhibited decreased reactivity which predicts a more closed hairpin structure in the absence of RBM42 (**Fig. 5b and Extended Data Figs. 6a,b**). To support the hypothesis that the remodeling of RNA Stem III with RBM42 loss may inhibit efficient translation, we constructed a luciferase reporter whose translation is driven by either the wildtype *MYC* 5’UTR or harbors the deletion of the newly formed hairpin. Critically, deletion of the hairpin residues significantly rescued the reduced translation efficiency observed with RBM42 depletion (**Fig. 5c and Extended Data Fig. 6c**). Next, we generated compensatory mutations that change the sequence but not the structure of the newly formed hairpin (**Extended Data Fig. 6a**). Mutation of these residues maintained the sensitivity of *MYC* 5’UTR dependent translation to RBM42, corroborating the findings that structure acts as a translation repressor (**Extended Data Figs. 6c,d**). Taken together, these data suggest that RBM42 can directly bind the *MYC* 5’UTR to remodel the RNA structure to promote more efficient translation initiation and maintain Myc protein levels.

As RBM42 also binds to the *EGFR* and *JUN* 5’UTRs to alter translation efficiency, we asked if there were common RNA structures that might be recognized and bound by RBM42. We employed DMS probing and targeted sequencing to determine the RNA structures of the *EGFR* and *JUN* 5’UTRs in PDAC cells (**Extended Data Figs. 6e,f and Supplementary Table 7**). While the analysis revealed that, like *MYC*, the *EGFR* and *JUN* 5’UTRs are highly structured, a comparison of the regions bound by RBM42 in each 5’UTR did not yield any striking structural similarities. Therefore, RBM42 directly binds a specific set of highly structured 5’UTRs to promote more efficient translation initiation to coordinate the expression of a pro-oncogenic program.

### RBM42 is a ribosome associated protein that interacts with the 43S pre-initiation complex

To better understand the mechanism by which RBM42 regulates translation, we performed cellular fractionation and immunoprecipitation (IP) of the endogenous cytoplasmic RBM42 in Panc-1 PDAC cells followed by unbiased mass spectrometry analysis. These experiments revealed that the cytoplasmic RBM42 protein-protein interactome was enriched for proteins with RNA binding functions and several components of the translation machinery (**Figs. 5d,e and Supplementary Table 8**). In particular, the RBM42 interactome contained the majority of small subunit (40S) ribosomal proteins (20/33), in addition to key translation initiation factors, such as eIF2 (eIF2α, β, and γ) and eIF4F members (**Fig. 5d**). By immunoprecipitation and western blot analysis, we confirmed cytoplasmic RBM42 binding to 40S components RPS12, RPS3, RPS20, and RPS8, as well as a to the eIF2 complex component eIF2β, which were not detected in the IP of nuclear RBM42 (**Fig. 5f and Extended Data Figs. 6g,h**). RNAse treatment did not alter RBM42 interaction with the 40S ribosomal proteins (RPS) and eIF2β, supporting direct interactions (**Fig. 5f**). The cytoplasmic interaction between RBM42 and RPS or eIF2β was further supported by proximity ligation assay experiments, which showed RBM42 near RPS and eIF2β in PDAC cells *in vivo* (**Fig. 5g**). Consistent with the IP mass spec data, we observed a low level interaction of RBM42 with eIF4E and eIF4A1, factors important for oncogene translation^10,11^, but not with eIF5A^29^ , nor with cytoplasmic proteins that were not present in the RBM42 interactome (**Extended Data Fig. 6i and Supplementary Table 8).**

The protein-protein interaction data suggest that RBM42 may interact with the 43S pre-initiation complex (43S PIC), which contains the 40S ribosomal subunit, initiation factors eIF1, eIF1A, eIF3, eIF5 and the eIF2-Met-tRNAi^Met^-GTP ternary complex^57^. The eIF4F cap-binding complex recruits the 43S PIC to the mRNA, forming the 48S PIC and begins scanning to find the start codon, functioning as a key rate limiting step for translation initiation and protein synthesis. In support of RBM42’s role in enhancing the 43S PIC recruitment and thereby 80S formation, polysome fractionation followed by western blotting showed that RBM42 was predominantly found in the initiating fractions (**Fig. 5h**). Treatment with EDTA that leads to the disassembly of 80S ribosomes resulted in the strong co-fractionation of RBM42 with the 40S (**Fig. 5h**). This polysome fractionation analysis strongly supports a role for RBM42 as a ribosome binding protein (RAP), which, in this capacity, may directly bind components of the pre-initiation complex. Thus, PDAC cells may co-opt RBM42 to enhance and direct the pre-initiation complex to key oncogenic mRNAs, such as *MYC*, thereby modulating their translation initiation efficiency and tuning their protein dosage.

To test this hypothesis, we employed the 48S PIC assembly assay to determine if RBM42 mechanistically functions to promote the recruitment of the 43S PIC to the *MYC* mRNA. In this assay^58^, Panc-1 cytoplasmic lysate is treated with non-hydrolyzable GTP (GMP-PNP) to lock the recruited 43S PIC onto the RNA, at which point it is referred to as the 48S PIC^57^ (**Fig. 5i**). PDAC cell lysates with control or RBM42 knockdown were treated with GMP-PNP and then subjected to sucrose gradient fractionation (10-30%) followed by qPCR to quantify the amount of 48S PIC assembled on the *MYC* RNA compared to the unbound RNA fractions. Depletion of RBM42 reduced the assembly of the 48S pre-initiation complex on the *MYC* mRNA, but did not impact recruitment to a control mRNA, *TUBB* (**Fig. 5j and Extended Data Fig. 6j**). These findings support a model wherein RBM42 regulates the recruitment and scanning of the pre-initiation ribosomal complexes on the *MYC* 5’UTR to sustain the necessary level of Myc protein production in PDAC cells.

### RBM42 regulates *in vivo* tumorigenesis through Myc

We sought to understand the role of RBM42 in PDAC tumorigenesis. Examination of publicly available proteomics data of human PDAC, showed that RBM42 and Myc protein levels are significantly correlated and show a strong trend in predicting patient survival (**Figs. 6a,b**). In contrast, *RBM42* and *MYC* mRNA expression are uncorrelated in the same patient cohort, supporting a translational control mechanism *in vivo* (**Fig. 6c**). To provide further clinical relevance, we examined RBM42 and Myc expression in TCGA RNA sequencing data and human PDAC tissue. First, we stratified PDAC TCGA data to separate tumors with high and low *RBM42* RNA expression and examined the enriched gene signatures in each cohort. Tumors with the highest *RBM42* expression displayed significant enrichment for “MYC target” gene signatures as compared to the lower *RBM42* expressing tumors (**Fig. 6d and Extended Data Fig. 7a**). These data demonstrate that tumors with the highest *RBM42* expression also possess high Myc transcriptional activity, which is indicative of high Myc protein levels. Moreover, we examined RBM42 and Myc protein expression co-occurrence in PDAC ductal structures in a panel of human PDAC tumors. The majority of ducts positive for Myc protein also had high RBM42 expression consistent with co-expression of these proteins and a role for RBM42 in regulation of Myc protein dosage (**Fig. 6e and Extended Data Fig. 7b**). Accordingly, most ducts with low RBM42 levels were also negative for Myc protein (**Extended Data Fig. 7c**). RBM42 was observed to localize in both nuclear and cytoplasmic compartments in PDAC tissue (**Fig. 6e and Extended Data Fig. 7b**). Together, these data indicate a strong correlation between high RBM42 and high levels of Myc protein expression, supporting a functional role of RBM42 in selective translational control *in vivo* in PDAC tumors.

**Figure. 6:**
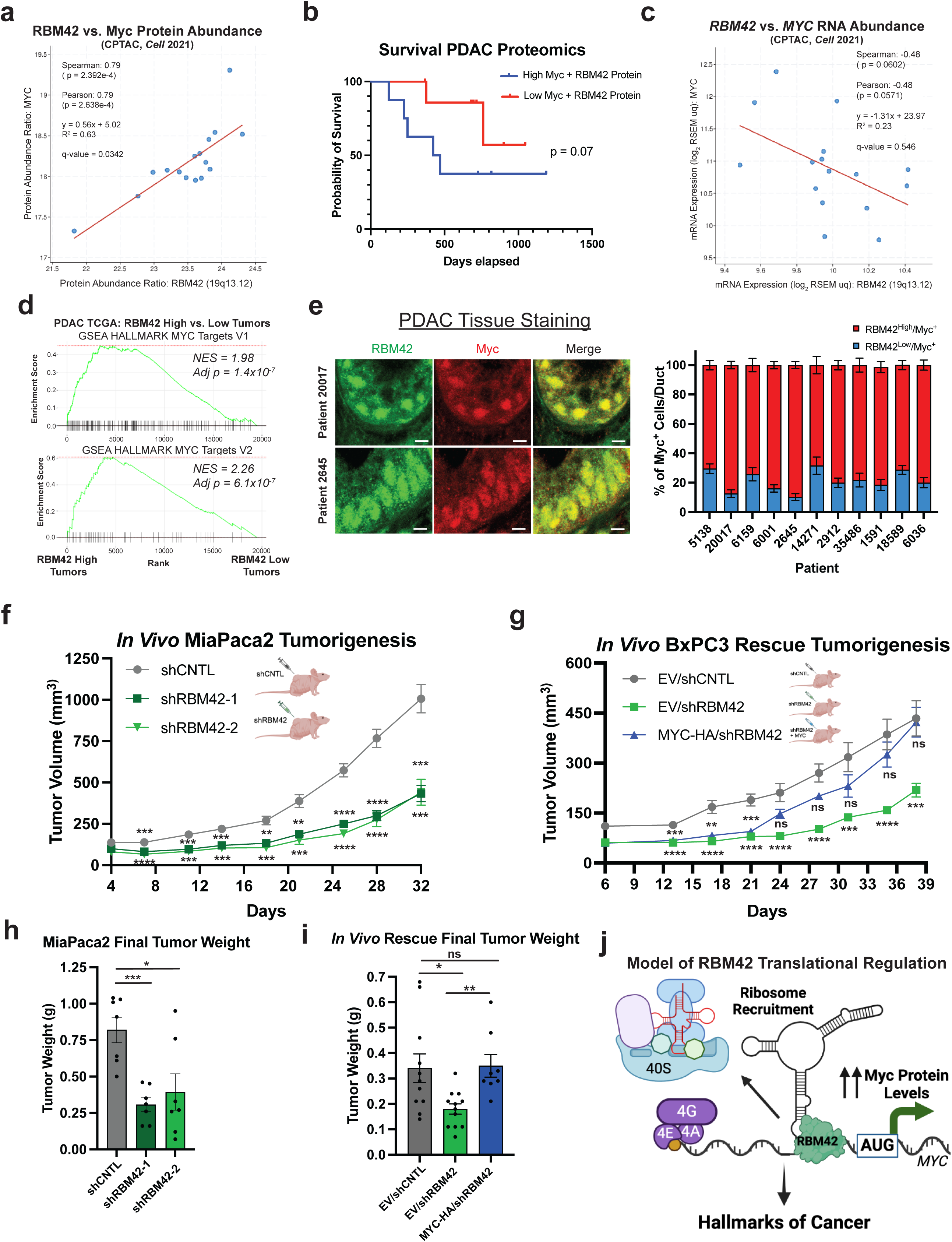
RBM42 regulates *in vivo* PDAC tumorigenesis. **(a)** Correlation of RBM42 and Myc protein abundance from human pancreatic adenocarcinoma specimens^45^, n=16. Graph and statistics generated with cBio Cancer Genomics Portal^67^. **(b)** Kaplan-Meier survival plot comparing patients in (a) with high Myc+RBM42 compared to low Myc+RBM42 protein levels. **(c)** Correlation of *RBM42* and *MYC* mRNA abundance in the same human pancreatic adenocarcinoma specimens as in (a). **(d)** MYC Targets gene set enrichment analysis (GSEA) comparing top quartile vs. bottom quartile of *RBM42* RNA expression in TGCA PAAD RNA-sequencing data (n=45 patients/quartile), Normalized Enrichment Score (NES). **(e)** Representative Myc and RBM42 immunofluorescence co-staining in human PDAC tissue and quantification of RBM42 and Myc per cell overlap/patient; n = 6-49 ducts/patient, mean ± SEM. Scale bar = 5μm. **(f)** Tumor volumes of MiaPaca2 pancreatic cancer cell xenografts with control and RBM42 knockdown, n=7 mice/group. **(g)** Tumor volumes of BxPC3 pancreatic cancer cell xenografts with control, RBM42 depletion or RBM42 depletion with enforced Myc expression (coding sequence), n= 8-12 mice/group. **(i)** Final tumor weights of BxPC3 xenografts. **(j)** Model of RBM42 selective translational control of pro-tumorigenic mRNAs, as exemplified by *MYC*. All graphs mean ± SEM unless noted. * p < 0.05, ** p < 0.01, *** p <0.001, **** p <0.0001 (Unpaired t-test).

Finally, we examined the necessity of RBM42 for pancreatic cancer tumor growth *in vivo* in two different tumor models. Loss of RBM42 expression resulted in diminished tumor growth and smaller final tumor size in both MiaPaca2 and BxPC3 PDAC xenograft models (**Figs. 6f-i and Extended Data Figs. 7d,e**). To determine to what extent decreased Myc protein levels are responsible for reduced tumorigenesis, we constitutively expressed the Myc coding sequence in PDAC cells and tested their ability to grow *in vivo* following RBM42 suppression. Enforced Myc expression was sufficient to rescue BxPC3 tumor growth *in vivo* (**Figs. 6g,i and Extended Data Fig. 7e**). These results strongly imply that, while RBM42 controls the translation of other oncogenic factors, a predominant function of RBM42 in tumorigenesis is regulating Myc protein expression. These data substantiate that increased RBM42 activity acts as a *bona fide* oncogenic lesion through its selective translational control of Myc expression in human PDAC (**Fig. 6j**).

## DISCUSSION

Here we use a functional, genome-wide approach to identify a regulatory network of transcript-specific *trans*-factors that promote *MYC* mRNA translation in PDAC and functionally characterize top hits. Supporting the richness and depth of this new data set, we used gold-standard polysome profiling to validate four new selective positive regulators of *MYC* translation whose functional activity in PDAC can be studied in the future. In particular, we characterized UBAP2L, a multifunctional protein posited to be associated with the ribosome, whose RNA is elevated in pancreatic cancer and protein levels increase with PDAC tumor stages^59^. Additionally, we demonstrated the selective regulation of *MYC* mRNA translation by the m^6^A “writer” METTL3 and “reader” YTHDF2, whose RNA expression levels are each predictive of negative prognosis in PDAC^48^. These data illustrate how cancer cells have adapted diverse means to maintain sufficient dosage of key oncogenic proteins.

Excitingly, we identified RBM42 as a top candidate activator of *MYC* translation. While RBM42 has been little studied, particularly in cancer, we found that its expression is increased in PDAC and is correlated with poor patient survival. Moreover, both UBAP2L and RBM42 protein expression levels were correlated with Myc protein levels in a panel of human PDAC tumors, suggesting the applicability of our newly identified selective translational control mechanisms *in vivo*. Our studies represent an important step towards fully unraveling the complexity of selective translational control of specific oncogenes as well as the coordinated protein synthesis of oncogenic programs. The unique repertoire of RBPs that selectively regulate the translation of driver oncogenes has remained elusive. However, this highly adaptable functional screening approach allows us to begin to map out how cancer cells tune the expression of oncogenes. Future studies that expand this approach to additional oncogenes and cancer types will reveal patterns of common and unique translational regulation, which will provide critical opportunities to identify new therapeutic targets.

We identified RBM42, as a key regulator of *MYC* mRNA translation and a suite of select, yet vital pro-oncogenic genes. Indeed, our *in vivo* rescue experiment suggests that, while Myc is a major translation target and oncogenic determinant downstream of RBM42, sustained expression of the entire coordinated RBM42 oncogenic translational program is required for the complete rescue of PDAC tumor growth dynamics. This work demonstrates the power of a multi-omics approach, synthesizing transcriptomic, translatome, proteomic, CLIP-seq and RNA structure determination methods, to uncover novel regulatory mechanisms governing the expression of dominant oncogenes like Myc. This is exemplified by our determination of the endogenous *MYC* 5’UTR RNA structure, for the first time, which illuminates the ability of RBM42 to remodel RNA structure and serve as a critical interface for the recruitment of the translation pre-initiation complex.

There is an increased appreciation of the diversity and importance of ribosome-associated proteins^60,61^, although their functional roles have remained elusive, particularly in cancer etiology. In this study, we discovered that RBM42 binds to ribosomes likely in the context of the pre-initiation complex to promote translation initiation of specific mRNAs. In particular, the RBM42 protein interactome prominently includes the eIF2 complex, which suggests RBM42 binding to the mRNA entry channel face of the 43S pre-initiation complex where eIF2 binds^57^. Moreover, our data support a direct interaction between RBM42 and eIF2β (Fig. 5f). eIF2β has been shown to enhance PIC assembly, scanning, and fidelity of start codon selection^62–64^. Therefore, RBM42 may influence translation initiation efficiency through its interaction with eIF2β on select target transcripts. Our work demonstrates that cancer cells harness RBM42 to selectively bind and alter the 5’UTRs of oncogenes to increase translation efficiency and increase protein dosage. We speculate that, with enhanced bioinformatic and experimental tools, shared 5’UTR RNA structural elements may be identified among the RBM42 translational target genes, representing an interesting future direction to explore.

A few previous studies have implicated RBM42 in response to cell stress states. In particular, a recent study implicated RBM42 in response to DNA damage with a narrow focus on its translational control of the *CDKN1A* mRNA^47^, which was previously shown to be bound by RBM42^65^. This is interesting as cancer cells are under persistent stress, such as aberrant proliferation, hypoxia, the harsh conditions of the metastatic process as well as chemotherapeutic treatments^12^. In this context, it is critical to emphasize that selective translation of pro-survival factors is a rapid response mechanism to dynamically reshape the cancer cell proteome, depending on the changes in the tumor environment. Myc is a central contributor to promoting cancer cell growth and adaptation to therapy, leading to drug resistance and tumor progression, particularly in PDAC^22,25,26^. Interestingly, it was reported that in the context of a synthetic reporter construct downstream of inducible Myc expression in lymphoma cells, RBM42 may regulate the translation of certain Myc target genes^46^. However, it is entirely unclear the endogenous relevance of these findings or any potential mechanism or impact on cancer cells. Given that RBM42 regulates a central pro-oncogenic translation program, it may be an essential coordinator of the metastatic process and response to chemotherapeutic treatments, helping cancer cells to persist under therapy.

Our mechanistic and *in vivo* data highlight RBM42 as a potent therapeutic target whose increased protein expression in cancer cells could provide a necessary therapeutic window. Our work deciphers a part of the translational code in cancer and offers a new opening to target the expression of deadly oncogenes in pancreas cancer. While genomic and transcriptomic analyses have laid the foundation for our understanding of cancer biology, selective translational regulators offer a new window into how the cancer proteome is ultimately shaped and expressed.

## ACKNOWLEDGMENTS

We would like to thank all members of the Ruggero laboratory, especially Duygu Kuzuoglu-Öztürk, for critical discussion of the project and for manuscript feedback. We thank Heather Karner (UCSF) for technical support for the CLIP experiments and Brian Zarnegar (Stanford University) for CLIP advice and irCLIP reagents. Sequencing was performed at the UCSF CAT, supported by UCSF PBBR, RRP IMIA, and NIH 1S10OD028511-01 grants. FACS and BC43 Spinning Disk Confocal microscopy was conducted in the HDFCCC Laboratory for Cell Analysis Shared Resource Facility through a grant from NIH (P30CA082103). Mass spectrometry was provided by the Mass Spectrometry Resource at UCSF (A.L. Burlingame, Director), supported by the Dr. Miriam and Sheldon G. Adelson Medical Research Foundation (AMRF). J.R.K. was funded by the American Cancer Society postdoctoral fellowship (133966-PF-20-009-01-RMC) and the UCSF Mentored Scientist Award in Pancreas Cancer (RAP 7031426). G.H. was supported by the National Science Foundation Graduate Research Fellowship Program (NSF2034836). H.G. is an Arc Institute Core Investigator. Research in the lab of Dr. Goodarzi is supported by the Arc Institute. R.M.P. is funded by R01CA240603 (NCI; NIH); AACR-MPM Oncology Charitable Foundation Transformative Cancer Research Grant; Ed Marra Passion to Win Fund. D.R. is funded by NIH (R35CA242986) and the American Cancer Society (American Cancer Society Research Professor Award).

## AUTHOR CONTRIBUTIONS

The study was designed by J.R.K. and D.R. J.R.K, G.S., M.S., N.M, and K.Y. performed experiments. V.S., I.L. and S.M. conducted the bioinformatic analyses with supervision and input from H.G.. G.H., G.R. and R.P. provided human and mouse pancreas cancer specimen sections, aided in tissue staining, imaging and quantification, and contributed conceptually to this project. K.W.W. and G.E.K. provided human pancreas cancer specimens. J.A.O.-P. and A.L.B. performed mass spectrometry analysis. J.R.K., G.S. and D.R. wrote the manuscript with review and editing from all other authors.

## METHODS

### Cell Lines

All cell lines were cultured at 37 °C, 5% CO2. Lenti X 293T, Panc-1, and MiaPaca2 were cultured on DMEM 10% FBS, 1% Pen-Strep; BxPC3 and AsPC1 were cultured in RMPI-1460 10% FBS, 1% Pen-Strep. HPDE cells were cultured in Keratinocyte SFM, + EGF + bovine pituitary extract (Invitrogen Cat#:17005042), 1% Pen-Strep.

Panc-1, MiaPaca2, BxPC3 and AsPC1 lines were purchased authenticated from ATCC. Lenti-X 293T cells were purchased from Takara. HPDE cells were a gift from Rushika Perera’s lab, who have confirmed their identity. Lenti X 293T, AspC1, HPDE and BxPC3 are derived from female individuals. Panc-1 and MiaPaca2 are derived from male individuals. All cell lines were regularly confirmed to be mycoplasma-free by the MycoAlert Mycoplasma Detection Kit (Lonza).

### Animals

6-week old, female Athymic nude mice (Hsd:Athymic Nude-Foxn1^nu^) were purchase from Invigo. Animal cohort numbers were chosen based on results from tumorigenesis pilot studies in mice. Animals were randomly assigned to experimental groups. Animals were not involved in any previous procedures prior to subcutaneous tumor injection. Researchers were not blinded to experimental group identity. Mice were housed five to a cage with autoclaved bedding, food, and water. All animals under assessment were weighed at least once a week to observe any changes indicative of poor health status. All the study procedures related to mouse handling, care, and assessment were performed according to the guidelines approved by the University of California San Francisco Institutional Animal Care & Use Committee (IACUC).

### Human Specimens

Human pancreatic cancer specimens were provided by the UCSF Department of Anatomy. The samples were collected under IRB approved protocols and are deidentified to ensure patient confidentiality.

### Plasmids and Viral Transduction

The fluorescent translational reporter was constructed beginning with the pLV-EF1a-IRES-Blast (Addgene #85133) with the hPGK-mCherry sequence from pLenti.PGK.chFP.W (Addgene #51008) was cloned in reverse orientation of the EF1a promoter using In-Fusion (Takara) cloning. The destabilization domain was added to the 3’ end using inverse PCR. Finally, the *MYC* 5’UTR (IDT gBlock) sequences and the destabilized GFP were cloned into the MluI site using multi-piece In-Fusion cloning to ensure seamless insertion. The destabilized GFP sequence was cloned from FUGW-d2GFP-ZEB1 3’UTR (Addgene #79601). All cloning was transformed into either lab made DH5α or Stellar (Takara) competent bacteria.

All exogenous expression constructs were cloned into LentiORF pLEX-MCS-IRES-Puro (Gift of P.A. Khavari lab). In-Fusion (Takara) was used to insert RBM42 or Myc-HA into the BamHI/XhoI sites. The human RBM42 (MHS6278-202830315) and Myc (MHS6278-202755482) cDNA sequences were obtained from Horizon Discovery Biosciences Ltd. In all cases, only the coding sequences (CDS) were PCR amplified from the commercially obtained cDNA sequences for cloning into the exogenous expression constructures. In the case of Myc, the sequence encoding the 439 amino acid isoform was used, beginning with the main ATG.

ShRNA knockdown was done with pLKO.1 (Addgene #10878) backbone. For pLKO.1 shRNA hairpin cloning, sense and anti-sense insert oligos were annealed and ligated into the cut backbone using T4 DNA ligase at room temperature for at least 1 hour. To validate candidate regulators, cells were infected with shRNAs. After 72 hours, the cells were harvested via trypsinization and split for protein and RNA expression analyses. Hairpin sequences are listed in Supplementary Table 9.

SgRNA were cloned into pLG vector (BFP-expressing, puromycin selectable; gift of Luke Gilbert lab) cut with BstX1 and BlpI. Sense and anti-sense oligos containing the target sequences with annealed in 1x annealing buffer (100mM Potassium acetate, 30mM HEPES-KOH pH 7.4, 4mM Magnesium acetate) 95C, 5min then cool to room temperature on bench. 1:20 diluted oligos were ligated (T4 ligase NEB) into 50ng of cut and purified vector then transformed into bacteria. Target sequences are listed in Supplementary Table 9.

For both exogenous expression, sgRNA and shRNA lentiviral production, 293T were transfected with 9μg lentiviral expression construct, 7μg psPAX2, and 2.5μg PMD2.G. Transfections were done in 10cm plates using Lipofectamine 2000 (Life Technologies). Viral supernatant was collected at 48 and 72 hours after transfection and concentrated using Lenti-X Concentrator (Takara). All cell lines were seeded into lentiviral-containing media and transduced overnight with 5μg/mL polybrene with fresh media change the next morning. Cells were selected using 1-5μg/mL puromycin or blasticidin, or hygromycin 80-150μg/mL approximately 36 hours after initial infection. To generate the inducible CRISPRi cells as well as the reporter cells, the cells were induced with doxycycline (5μg/mL) for 3 days were sorted to isolate the brightest cell population.

### Immunoblotting

For protein extraction, cells were lysed in 1X RIPA (50mM Tris pH 8.0, 250mM NaCl, 1% IGEPAL CA-630, 1mM EDTA, 1% Sodium Deoxycholate, 0.1% SDS) with Complete Mini Protease Inhibitor Cocktail, EDTA-free (Millipore Sigma) and PhosStop phosphatase inhibitor cocktail (Millipore Sigma). Cells were allowed to lyse on ice for at least 15 minutes, sonicated if needed, and centrifuged at 16,000xg for 15 minutes. The soluble fraction was removed and quantified using the Pierce BCA Protein Assay Kit. 15-30ug of protein was loaded into Mini Protean TGX 4-15% gels (BioRad) and transferred to 0.45μM pore size Nitrocellulose. Membranes were blocked with Odyssey blocking buffer (PBS) (LI-COR) for at least 30 minutes at room temperature. Primaries were added to 5% BSA in TBST and incubated with the membrane overnight at 4C. Loading control antibodies-β-Actin and Gapdh-were incubated for 1hr at room temperature. The membranes were washed three times in TBST and incubated with LI-COR secondary antibodies-800CW Goat anti-Mouse IgG (H + L), 800CW Goat anti-Rabbit IgG (H + L), 680RD Goat anti-Mouse IgG (H + L)-(1:15,000) in 5% milk in TBST for 45 minutes at room temperature. The membranes were washed twice in TBST and once in PBS then scanned on the Odyssey CLx. All quantification was done in the Image Studio Lite software.

The following antibodies were used: anti-β-Actin (A1978, Millipore Sigma), anti-UBAP2L (A300-533A, Bethyl), anti-METTL3 (A301-567A, Bethyl), anti-YTHDF2 (24744-1-AP, ProteinTech), anti-c-Myc (D84C12) (5605S, Cell Signaling), anti-RBM42 (A305-139A, Bethyl), anti-EGFR (4267, Cell Signaling), anti-c-Jun (Cell Signaling, 9165), anti-SRF (Cell Signaling, 5147), anti-PAK4 (Bethyl, A300-356A), anti-EZR (Cell Signaling, 3145), anti-RPS6 (54D2) (Cell Signaling, 2317S), anti-RPL23 (Bethyl, A305-008A), anti-puromycin (Millipore, MABE343), anti-GAPDH (Cell Signaling, 5174S), anti-Lamin A/C (Cell Signaling, 4777S), anti-LSD1 (Cell Signaling, 4064S), anti-α-tubulin (Sigma, T8203), anti-EIF2S2 (Bethyl, A301-743A), anti-RPS12 (ProteinTech, 16490-1-AP), anti-RPS3 (Cell Signaling, 9538S), anti-RPS20 (ProteinTech, 15692-1-AP), anti-RPS8 (Bethyl, A305-016A), anti-HA epitope tag (HA.11) (BioLegend, 901513).

### Quantitative polymerase chain reaction (qPCR)

RNA from cell lysates was isolated using RNeasy Plus (Qiagen) according to manufacturer’s instructions. Single-stranded cDNA was synthesized by using 0.5-1μg RNA with High-Capacity cDNA Reverse transcription kit (Applied Biosystems). cDNA samples were diluted 1:20 and 1-2 μL of template was used in a PowerUP SYBR Green master mix reaction run on an QuantStudio 6 Flex Real-Time PCR System (Applied Biosystems). qPCR primer sequences are listed in Supplementary Table 9.

### Selective Translation Reporter CRISPRi Screen

#### CRISPRi Screen

For the CRISPRi screen, sgRNA library virus was generated from a commercially available human top 5 guide whole genome library (Addgene #83969). This virus was titrated on the Panc-1 CRISPRi 5’UTR Myc reporter cell line ensure 1 guide per cell. With the appropriate amount of virus, cells were infect for 1000x coverage of library. The next day media was changed and the subsequent day puromycin selection began to ensure every cell expresses an sgRNA. After 2 days of puromycin selection, when the negative, uninfected controls were dead, the media was changed to regular media. After this 24 hour recovery period, the cells were checked for BFP positivity to quantify infection percent (target is 20-50% positivity). Simultaneously, 1000x coverage cells were seeded into doxycycline (5μg/mL) containing media in 15cm plates. Doxycycline was refreshed daily and the cells were split as needed, always maintaining 1000x coverage. No cell stress or cytotoxicity was observed with the addition of doxycycline.

On day 5 of doxycycline treatment, working in batches, the cells were trypsinized and prepared into a single cell suspension for sorting. For sorting, the cells were first gated on BFP^+^ cells (cells with sgRNA) and then the mCherry negative cells were gated out. Finally, gates were set to collect the top 25% and bottom 25% on the GFP/mCherry ratio. The cells were sorted until at least 25,000,000 for each top and bottom 25%, that is at least 50,000,000 cells total, using an BD Aria 3 sorter. This was repeated for two replicates (independent sgRNA library infections). Cell pellets were snap frozen for library preparation.

#### Library Preparation, Sequencing and Analysis

For library preparation, genomic DNA (gDNA) was isolated independently for the top and bottom fractions for both replicates (4 samples total) using the Macherey Nagel Nucleospin Blood XL kit. The gDNA per sample was quantified and the same amount of gDNA was used for each pair of samples for the PCR amplification and Illumina barcoding. The following primers were used Forward:aatgatacggcgaccaccgagatctacacgatcggaagagcacacgtctgaactccagtcacXXXXXXgcacaa aaggaaactcaccct (X = index)

Universal Reverse: CAAGCAGAAGACGGCATACGAGATCGACTCGGTGCCACTTTTTC with the following conditions: 1x 98C 30sec; 22x 98C 10sec, 65C 75sec; 65C 5min Q5 MasterMix (NEB). All PCR reactions were pooled and 10% of the total reactions were cleaned up with SPRI Select beads with two-sided selection for a 270bp target product size. The libraries were quantified and confirmed size using BioAnalyzer HS DNA analysis. Libraries were pooled in equimolar ratios. Libraries were sequenced on a HiSeq400 with custom Sequencing primer: GTGTGTTTTGAGACTATAAGTATCCCTTGGAGAACCACCTTGTTG with 10% Phi-X spike-in.

The resulting data were analyzed with a publicly available state of the art workflow (https://github.com/mhorlbeck/ScreenProcessing)^68^. Briefly, the analysis pipeline maps the sgRNAs to the library of sgRNAs and counts the sgRNAs in the raw sequencing files. Then, it calculates the sgRNA-level and gene-level phenotypes and p-values. Mann-Whitney p-values are calculated by comparing the phenotypes of all gene targeting sgRNAs to the negative control sgRNAs included within the library. For this screen, the top and bottom pools were compared to generate the phenotype scores. There was no normalization for doubling time as the initial population was the same before sorting. The sequencing detected 18,900 out of the 18,905 total sgRNAs. First, the hits were filtered by discrimination score (DS) DS=average phenotype score*-log_10_[Mann-Whitney p-value] < -1.0 or > 1.0. To refine the hits further, we set a threshold for average phenotype score > 0.75 or < -0.75 and a p-value < 0.05. This analysis resulted in 622 candidate activators and 309 candidate repressors of *MYC* translation initiation.

### Single Guide Validation Studies

Based on the sequencing analyses, the top 2 sgRNAs were chosen for candidate regulators. The single sgRNA were cloned as described above. Panc-1 CRISPRi 5’UTR *MYC* reporter cells were transduced with the guides and puromycin selected. These cells were then seeded into 12-well plates subjected to 5 days of doxycycline treatment and analyzed using flow cytometry. Matched cells were also harvested to check for RNA depletion of the target gene. Flow Jo was used to gate and quantify the mean fluorescence intensity of GFP and mCherry.

### Sucrose Gradient Fractionation and Polysome Profiling

Panc-1 cells with control or targeted knockdown (72hrs post-infection) at about 60-80% confluency were then treated with 0.1 mg/mL cycloheximide for 3 min, scraped into PBS with 0.1mg/mL cyclohexamide, all plates were pooled together and centrifuged 3mins 300xg. Cell pellets were lysed in 250uL of lysis buffer (20 mM Tris pH 7.5, 150 mM NaCl, 5 mM MgCl_2_, 1 mM DTT, 1% Triton X-100, 0.1 mg/ml cycloheximide, 200 U/ml RNasin (Promega)). Nuclei and membrane debris were then removed by centrifuging at 15,000x g, 10 min. Equal amounts of protein was loaded onto a sucrose gradient (10%–50% sucrose(w/v) made with Gradient Master™ (BIOCOMP), 25 mM Tris pH7.5, 25 mM NaCl, 5 mM MgCl_2_, 1mM DTT) and centrifuged in a SW40 rotor (Beckman) for 2.5 h at 38,000 rpm at 4°C. 14 fractions were collected by density gradient fractionation system Piston Gradient Fractionator™ (BIOCOMP).

For RNA distribution analysis, RNA was isolated from fractions 3-14 using Trizol-chloroform followed by isopropanol precipitation >2hrs at -80C, 70% ethanol wash and resuspended in water. qPCR data were normalized to an FLuc RNA spike in prior to RNA preparation. Statistical analyses were done using 2-way ANOVA with uncorrected Fisher’s LSD test.

For western blot analysis, the ProteoExtract Protein Precipitation Kit (Sigma Millipore) was used with the following modifications. Prepared precipitant was added at an equal volume to the sucrose gradient fraction and left at -20C for at least 48hrs. The samples were centrifuged 10mins 16,000 xg at room temperature, the supernatant was removed, and the pellet was washed with 0.75mL ice cold prepared wash solution and centrifuged 5min. The supernatant was removed, and the pellet was again washed with 0.5mL wash solution, centrifuged 5mins and all remaining liquid removed. The pellet was allowed to dry, resuspended in 1x laemmli sample buffer, 5% BME and 1x RIPA at 80C, 10mins, 1000 rpm constant shaking.

### Puromycin incorporation assay

Cells were infected with control (GFP) or RBM42 targeting hairpins (blasticidin selectable pLKO) 72hours prior to the puromycin incorporation assay. For the cycloheximide control, cells were pre-treated with 50μg/mL cycloheximide for 15 mins. All conditions were treated with 1μM puromycin for 30 mins. Cells were then harvested via trypsinization and snap frozen for western blot analysis.

HPDE overexpression experiments

HPDE cells were infected overnight (∼16hrs) with 0.5-1x concentrated pLEX empty vector (EV) control or RBM42 expressing lentivirus in 6cm plate. The next morning, old media with the virus was removed, and the cells were moved to a 10cm tissue culture plate. 72 hours post-infection, the cells were harvested for protein and RNA analyses. For the enforced Myc expression experiments, HPDE cells were infected as described and then selected with puromycin to ensure consistent and stable Myc expression. After selection, cells were replated at different days for independent replicates of RNA and protein expression.

### Cycloheximide Protein Stability Assay

Cells infected with either control or RBM42 knockdown were seeded into 6-well plates. 72hours after shRNA infection, cells were treated with 10μg/mL cycloheximide for 0, 15, 30, 45, 60 and 90mins. Cells were harvested with cold PBS and scraping. The stability of Myc protein and the knockdown were assessed with western blotting. Both bands were used to quantify Myc protein levels.

### Cell Phenotyping Experiments

PDAC cells either parental or for rescue experiments, previously generated, hygromycin empty vector (EV) or Myc-HA expressing cells, were infected with control or RBM42 knockdown hairpins. The next day media was changed. The following day (∼36hrs post-infection) the cells were placed in the puromycin selection. After 2 days of puromycin selection, the cells were allowed to recover in standard media for 24 hours. The same cell population was then seeded into 12-well plates (0.15-0.4×10^5^ cells/well) for cell titer glo timecourse (d0-d4), at extremely low confluency (0.02-0.2×10^5^ cells/well) in 6-well plate format for 2D colony formation or layered into soft agar. The remaining cells were split for RNA and protein analysis of knockdown. The Cell-titer Glo Assay (Promega) was performed following the manufacturer’s protocol and the absorbance at 490nm was measured on Glomax plate reader (Promega).

For counting the total live cells across the timecourse, cells were seeded as described for the cell titer glo experiments. At each time point, a plate of cells was trypsinized and quenched in a constant volume. An aliquot of cells was mixed with trypan blue and live cells were counted. Total cell numbers were normalized to the number of cells seeded.

For the 2D colony formation, after approximately 1.5 to 3 weeks, depending on visible colony size, the plates were fixed with ice cold methanol and then stained with 0.5% Crystal Violet (in 25% MeOH). Plates were washed with abundant water, imaged, and quantified using ImageJ (Fiji).

For the soft agar experiments, a 0.6% agarose dissolved in cell media base layer was set down in a 6well plate. Then, 0.05×10^5^ cells in a single cell suspension were mixed for 0.3% agarose final solution and allowed to solidify at room temperature before moving to a standard cell incubator. Cells were fed 1-2 times per week with a few drops of fresh media. After 4-6 weeks of growth, colonies were stained with 5mg/mL solution of MTT for ∼2hrs and imaged for manual quantification.

### Cytoplasmic/Nuclear Fractionation

Cells were harvested by scraping into ice cold PBS and pelleted 3mins 300xg. Cells were resuspended in 3-5 cell volumes of Buffer A- (10mM HEPES pH 7.4, 10mM KCl, 1.5mM MgCl_2_) and incubated on ice 5mins. An equal volume of Buffer A+ (10mM HEPES pH 7.4, 10mM KCl, 1.5mM MgCl_2_, 0.4% IGEPAL CA-630) was added and left to incubate on ice 2mins with periodic gentle mixing. The lysate was centrifuged in a refrigerated centrifuge at 4000rpm 1min and the cytoplasmic fraction was removed. No RNase treatment was performed. For nuclear fraction analysis, the remnant pellet was washed 1x in Buffer A-followed by lysis in 3x volume nuclei lysis buffer (50mM Tris pH 7.5, 0.05% IGEPAL CA-630, 10% glycerol, 2mM MgCl_2_, 250mM NaCl) for at least 10mins on ice. The nuclear lysate was cleared via centrifugation max speed, 10min at 4C. The lysate was removed as the nuclear fraction and the remaining pellet was discarded. Each protein fraction was spiked with 1x final Complete Mini Protease Inhibitor Cocktail, EDTA-free (Millipore Sigma).

### RBM42 RNA Expression Survival Analysis

For the analysis of the Yang et al. data^49^, median expression was used as the cutoff and Prism was used for the statistical analysis of overall survival. For the disease free survival analysis of Posta et al.^48^, the expression values for the top 3 tumor grades were used (G2, G3 and G4) and the analysis was done within KM plotter.

### Cytoplasmic CLIP Sequencing

Panc-1 cells were plated for ∼80% at the time of harvest (12x 15cm plates per replicate). One the day of harvest, media was changed and ∼3 hours later the media was removed. The plates were washed 1x with ice cold PBS. All PBS was thoroughly removed, the plates were placed on ice and crosslinked 400mJ/cm2 254nm. After crosslinking, the cells were scraped into ice cold PBS and cytoplasmic/nuclear fractionation was carried out as described above. Cytoplasmic lysate for each replicate was pooled and sonicated 2× 10sec at 20% amplitude (Branson Sonifier). The cytoplasmic lysate (at 3mg/mL) was digested with RnaseI (Ambion) diluted 1:5000 for 5min, 37C, mixing 15s 1000rpm and 45sec rest. Digestion was stopped with the addition of 2x volumes of cold lysis buffer. RBM42 was precipitated for 4.5hrs rotating at 4C with 1mg of protein with ∼2.3ug of antibody per IP pre-coupled to Protein G dynabeads.

After IP, the antibody-protein complexes were wash 1x High stringency (20mM Tris pH7.5, 5mM EDTA, 1% Triton X-100, 1% Sodium deoxycholate [Na-DOC], 0.001% SDS, 120mM NaCl, 25mM KCl) 5mins, 1x High salt (20mM Tris pH7.5, 5mM EDTA, 1% Triton X-100, 1% Na-DOC, 0.001% SDS, 1M NaCl) 5mins and 2x quick NT2 (50mM Tris pH7.5, 150mM NaCl, 1mM MgCl_2_, 0.0005% Igepal) washes. CLIP-sequencing libraries were prepared using a previously published protocol^69^. Briefly, protein-bound RNAs were dephosphorylated on bead using T4 PNK (NEB) with modified PNK Buffer (10mM Tris pH 6.5, 50mM MgCl_2_, 50mM DTT) followed by PolyA tailing with yeast poly(A) polymerase (Jena). This was followed by 3′-end labelled with azide-dUTP, using yeast poly(A) polymerase (Jena) and labelled with IRDye800-DBCO conjugates (LiCor). The protein–RNA complexes were then eluted from beads, resolved on a 4–15% gel, transferred to nitrocellulose and imaged using an Odyssey CLx instrument (LiCor). Regions of interest were excised from the membrane, and the RNA was isolated by Proteinase K digestion and oligo-dT dynabead purified in high salt (500mM NaCl) buffer to promote capture. RNA was eluted into TE buffer (20mM Tris pH 7.5, 1mM EDTA) followed by library preparation using SMARTer smRNA-Seq Kit (Takara), beginning with reverse transcription (custom RT primer:CAAGCAGAAGACGGCATACGAGATNNNNNNNNGTGACTGGAGTTCAGACGTGTGCTCTTCCGATCTTTTTTTTTTTTTTT, Ns = UMI). The library PCR, 20 cycles, was performed with index forward (i5) primers and universal reverse (P7) primer (CAAGCAGAAGACGGCATACGAG). The libraries were purified using SPRI select beads and sequenced as PE150 on Illumina NovaSeqX instrument at UCSF Center for Advanced Technologies.

To process the CLIP-Seq, adaptor sequences were first trimmed using *Cutadapt*^70^. Reads were then aligned to the human genome (build hg38) using bwa and the following parameters: *-n 0.06 -q 20*. PCR duplicates were collapsed from sorted bam files using *umitools*^71^. Finally, Peak calling and CIMS calling was performed using the *CTK* package^56^. Bed file outputs of clip peaks were then annotated to genomic features using *bedtools intersect*^72^.

For RNA linker ligation test, the RnaseI treatment was carried out as described above but using 1:15k, 1:5k, 1:1667, 1:555 titration. After PNK 3’ dephosphorylation, the ir800 labeled RNA linker was ligated on using RNA Ligase I (NEB), and 0.5uM of pre-adenylated adapter as previously published^73^. Ligation proceeded overnight at 16C, 15s 1350 rpm, 90s rest. The following day the ligation mixture was washed once with NT2, eluted from the beads and run on a gel as described above.

For RBM42 CLIP-qPCR experiments, the lysates were not treated with RNaseI. For cytoplasmic lysate ∼1.3mg of lysate was added to 2μg of rabbit IgG or RBM42 antibody coupled to Protein G dynabeads. For nuclear lysate, ∼0.3mg of lysate was added to 3μg of rabbit IgG or RBM42 antibody coupled to Protein G dynabeads. These ratios were based on titrations to capture >90% of RBM42 protein in each fraction. IP and washing was carried out as described above. After the final wash the beads with bound material were subjected to proteinase K treatment-380uL proteinase K buffer (100mM Tris pH 7.4, 100mM NaCl, 1mM EDTA and 0.5% SDS) + 20uL of proteinase K (Invitrogen AM2546)- for 45mins, 55C, constant shaking at 800rpm. The buffer was removed from the beads and combined with 400uL acid phenol:chloroform and incubated for 20mins 37C, constant shaking at 1000rpm. The aqueous layer was purified using a Heavy Phase Lock tube with one repeat purification with chloroform. The aqueous layer was removed and combined with 40uL sodium acetate, 1uL glycoblue precipitant and 500uL isopropanol and left to precipitate overnight at -80C. The next day the RNA was precipitated via centrifugation at 21,000 xg for 30mins at 4C, washed once with 70% ethanol, and resuspended in 20uL of water. 10uL of the resuspended RNA was used for cDNA synthesis with random primers.

### Polysome-Sequencing

Panc-1 cells with control or RBM42 knockdown in biological triplicate were subject to sucrose gradient fraction as described above. For each sample, equal volumes per fraction were pooled as follows fractions 3-6 for untranslated, fractions 7-10 for low translation and fractions 11-14 for high translation. A previously published polysome-sequencing protocol^74^ was followed with the following specifics. The RNA from each pool was purified using Trizol-chloroform followed by isopropanol precipitation. The RNA was resuspended in 20μl water and quantified using Qubit. Then 1 or 2μl of 1:200 ERCC RNA Spike-In Mix (Invitrogen, 4456740) was added to each fraction pool, depending upon the RNA amount. DNase treatment was performed with TURBO DNase (Ambion, AM2238) for 30□min at 37°C and samples underwent acid phenol extraction and were resuspended in 15μl of Ultra Pure DNase/RNase-Free Distilled Water. RNA concentration was quantified and 10ng was used as input into the SMARTer Stranded Total RNA-Seq Kit v3-Pico (Takara). Libraries from the total input RNA, 10ng, for each sample were prepared in parallel. Libraries were quantified for concentration and size distribution with BioAnalyzer HS DNA kit. They were sequenced PE150 on a NovaSeqX.

To analyze the polysome-sequencing data, similar to previously published methods^74^, Gencode V28 transcripts and ERCC control RNAs were separately quantified in the paired end samples using *salmon quant* (default parameters)^75^. Outputs were loaded using tximport^76^ and counts were normalized to the ERRC in the untranslated fraction (pooled fractions 3-6) within their respective samples. In particular, the lm() function in R was used to model the slope of the low translated (7-10) and high translated (11-14) fractions against the untranslated fraction (low/high ∼ untranslated). The slope of the line of best fit was then used to scale and normalize the low/high fractions to the untranslated fraction. These normalized counts were then input into EdgeR^77^ for differential expression analysis. We employed a design matrix that directly incorporated the interaction between fraction and condition (∼0 + fraction:condition). This approach allows us to analyze changes in the distribution of transcripts across the polysome profile without relying on pre-calculated ratios. The difference in translational efficiency (TE) between conditions was evaluated by comparing the relative abundance of transcripts in the highly translated fractions to those in the untranslated and lowly translated fractions. Specifically, we defined TE as [high translation] / [untranslated + low translation], but rather than calculating this ratio directly, we used the interaction terms in our model to assess how this relationship changes between conditions. This method captures shifts in translational status while accounting for sample-specific variances. Significant TE changes were called at adj p-value < 0.05 and FC < 0.75 or FC > 1.4)

### Total RNA-Sequencing Analysis

Salmon quant with default parameters was used to quantify paired end RNA-Seq reads. These reads were imported into DESeq2^78^ using tximport, and pairwise comparisons were performed using default DESeq2 parameters.

### Gene Set Enrichment Analyses

For the RBM42 regulated gene signature analysis, the fgsea library^79^ was used to perform gene set enrichment on DESeq2 outputs from the paired-end total RNA sequencing. Specifically, genes were ranked according to the *stat* ouput of DESeq2 differential expression tables. Then, the fgsea() function, using default parameters, was used to calculate the enrichment of hallmark pathways or the transcription factor targets msigdb collections^80^.

For the analyses of the TCGA pancreatic tumor gene expression, TPM-normalized RNA-Seq count matrices for 177 PDAC tumors from The Cancer Genome Atlas (TCGA) Pan-cancer dataset were retrieved using the cBio Cancer Genomics Portal^67^. Samples were ranked based on z-scored expression of RBM42 and participants in the upper and lower quartile of expression were selected. RSEM counts were input into DESeq2 and differential expression between high and low RBM42 expression groups was performed using default parameters. GSEA was performed as mentioned above.

### Global Splicing Analysis

To evaluate splicing changes in response to RBM42 knockdown, we used the rMATS-turbo pipeline^81^. Briefly, for triplicate samples per condition paired end total RNA-Seq reads were aligned to the transcriptome using STAR^82^ and splicing changes for exon skipping (ES), mutually exclusive exons (MXE), alternative 3’/5’ splice sites (A3SS/A5SS), and intron retention (RI) were statistically evaluated, using reads spanning splicing junctions. Percent spliced in (PSI) and FDR values for the respective event types were used for downstream visualizations and comparisons as noted in the figure legends.

### DMS-Seq protocol

Panc-1 cells were infected with hairpins 72hrs prior to Dimethyl Sulfide (DMS) treatment. Cells were harvested via trypsinization and split into untreated and treated aliquots. For DMS treatment, cells were resuspended in 2.8mL warm cell culture media, followed by 0.8mL 1M bicine buffer (pH 8.57 at room temperature). DMS was added to 1.6% final concentration and incubated at 37C for 5mins. The reaction was quenched with ice cold 2mL BME and cells were pelleted. Cells were washed once with 30% BME in PBS and RNA was purified using Trizol-chloroform as described above. To remove any contaminating genomic DNA, the RNA was treated with TurboDNAse per the manufacturer’s instructions and repurified. Reverse transcription of the endogenous MYC 5’UTR was done with Induro Reverse Transcriptase (NEB) and a reverse primer targeting the coding sequence proximal to the ATG. The cDNA was purified was SPRI select beads (2x volume). The cDNA was amplified using 5’UTR Myc specific forward and reverse primers using PrimeStar Max (Takara) for 35 cycles using ∼20% of purified cDNA product. The resulting DNA product was gel purified. Sequencing libraries were generated using the NEBNext Ultra II Kit following the manufacturer’s instructions with the following parameters: the input DNA was fragmented (FS DNA Module, NEB) for 40mins at 37C and 8 cycles for the library construction PCR. Product size and concentration were confirmed with BioAnalyzer HS DNA analysis. Samples were sequenced as PE150 on a NovaSeqX instrument. Primers for the reverse transcription and PCR amplification are in Supplementary Table 9.

DMS reads were trimmed using *trim galore* (default parameters) (https://github.com/FelixKrueger/TrimGalore), and then aligned to the reference *MYC* 5’UTR sequence using *bowtie2*^83^, using the parameters *-local --no-unal --no-discordant --no-mixed -X 1000 -L 12*. Bam files were then input into the perl-based *rnaframework* toolkit^84^. Mutations were counted using *rf-count -m* and normalized using *rf-norm (*parameters *-sm 4 -nm 2 -rb AC).* The region bound by the forward PCR primer was masked. Normalized replicate xml files were combined using rf-combine and structural prediction was done using rf-fold with parameters *-g - ct --folding-method 2 -dp.* The outputted ct files were imported into VARNA for visualization^85^.

### Luciferase reporter assay

The Myc 5′UTRs (WT and 363-394bp deletion mutant) were synthesized by IDT (gBlocks) and cloned into the pGL3 luciferase reporter vector (Promega) upstream of the Firefly luciferase open reading frame. Panc-1 cells with control or RBM42 knockdown were seeded into 6-well plates. Approximately 48 hours after initial infection, cells were transfected using Lipofectamine 3000 (Invitrogen) with 900ng of pGL3-FLuc-Sv40 and 100ng of RLuc. Media was changed ∼16hours later. 24hrs post-transfection cells were harvested: a fraction was saved for RNA extraction and the rest were lysed in a 100uL passive lysis buffer for at least 20 min on ice. The Rluc/Fluc activity was assessed using the Dual-luciferase Reporter Assay System (Promega) according to the manufacturer’s instructions, using a Glomax microplate luminometer (Promega). RNA extraction was performed using RNeasy (including gDNA elimination) kit following manufacturer’s instructions. Primers specific to FLuc and RLuc were spiked into cDNA to avoid any plasmid DNA contamination.

### Cytoplasmic RBM42 Immunoprecipitation Mass Spectrometry

For each replicate 10x 15cm of Panc-1 were seeded such that they were at 70-80% confluency at harvest with a complete media change 3 hours prior to harvest. Cells were harvested via scraping into ice cold PBS and spun down 3mins 300xg. Cytoplasmic and nuclear fractionation was carried out as described above. For each IP (IgG or RBM42), 3.5mg of cytoplasmic extract was combined with 7.5ug of antibody bound to Protein G Dynabeads and rotated overnight at 4C. The next day, the beads were washed 4x with wash buffer (50mM Tris pH 7.5, 0.1% IGEPAL CA-630, 5% glycerol, 50mM NaCl) followed by 3x PBS washes and 5x wash in water. Dry beads were snap frozen until preparation for mass spectrometry.

Sample-incubated streptavidin magnetic beads were resuspended in 9.4 ul of 5 mM Tris(2-carboxyethyl)phosphine (TCEP) 20mM triethylammonium bicarbonate (TEAB) pH 8.5, and incubated for 30 min at room temperature. After this, iodoacetamide was added to a final concentration of 7.5 mM and samples incubated for 30 additional minutes. 0.7ug of LysC (Wako) was added to each sample and incubated at 37C overnight. After that, 1 ug of MS grade trypsin (Thermo Scientific) was added to each sample and incubated at 37C for 4 hours. Supernatants of the beads were recovered into new low protein binding tubes and acidified with 5% formic acid. Peptides were recovered by solid phase extraction using C18 ZipTips (Millipore) and resuspended in 0.1% formic acid for analysis by LC-MS/MS. Peptide digests were subjected to chromatographic separation using a A 15-cm EasySpray C18 column (Thermo Scientific). 60 min water-acetonitrile gradients (2–25% in 0.1% formic acid) were used to separate peptides, at a flow rate of 200 nl/min, for analysis in a Orbitrap Exploris 480 (Thermo Scientific) in positive ion mode. MS spectra were acquired between 375 and 1400 m/z with a resolution of 120000. For each MS spectrum, multiply charged ions over the selected threshold (2E4) were selected for MS/MS in cycles of 3 seconds with an isolation window of 1.6 m/z. Precursor ions with a charge state of 2+ or higher were fragmented by HCD using a relative collision energy of 30. MS/MS spectra were acquired in centroid mode with resolution 30000 from m/z=120. A dynamic exclusion window was applied which prevented the same m/z from being selected for 30s after its acquisition. Peak lists were generated using PAVA in-house software^86^. All generated peak lists were searched against the human subset of the SwissProt database (SwissProt.2019.07.31) using Protein Prospector^87^. The database search was performed with the following parameters: a mass tolerance of 10 ppm for precursor masses; 30 ppm for MS/MS, cysteine carbamidomethylation as a fixed modification and acetylation of the N terminus of the protein, pyroglutamate formation from N terminal glutamine, and oxidation of methionine as variable modifications. All spectra identified as matches to peptides of a given protein were reported, and the number of spectra (Peptide Spectral Matches, PSMs) used for label free quantitation of protein abundance in the samples.

To confirm the interactions in a more stringent system, we expressed a c-terminally HA tagged eGFP or RBM42 in Panc-1 cells at approximately endogenous expression level. As described above, the cells were then subjected to cytoplasmic/nuclear fractionation. For RNase treatment, cytoplasmic lysate was subjected to RNaseA at 1μg/mL for 1 hour at 4C with constant rotation and stopped with 1μL/100μL RiboLoc. RNase digestion was confirmed through total RNA analysis following extraction with Trizol-chloroform followed by isopropanol precipitation. For cytoplasmic IP 1.25-1.6mg of cytoplasmic lysate or 75% nuclear lysate (to account for higher RBM42-HA concentration in the nuclear fraction) was combined with 30-40uL anti-HA magnetic beads (Pierce) for overnight IP. After IP, the beads bound with protein were washed 3x 3 minutes in cold wash buffer [50mM Tris pH 7.5, 0.1% IGEPAL CA-630, 5% glycerol, 50mM NaCl; 0.25% IGEPAL CA-630 was used for nuclear fraction to reduce background binding] and eluted in 2x laemmli sample buffer, 10 mins, 95C 1000rpm.

### 48S-PIC Assay

Panc-1 cells were infected with control or shRBM42 hairpins 72 hours prior to the assay. On the day of the experiment, the media as changed 3 hours prior to harvest. At harvest, the media was removed, and the cells were scraped into ice cold PBS with 0.1mg/mL cycloheximide. This protocol has been previously published^58^. The cells were pelleted 3mins 300xg and then lysed in 200-250uL RNL lysis buffer (10mM Tris pH 8.0, 140 mM NaCl, 1.5mM MgCl_2_, 0.25% NP-40, 0.15mg/mL cycloheximide, 20mM DTT) plus 40U/100uL of Rnasin Plus (Promega) 45mins on ice. The samples were further lysed with 5x through a 26G needle and then centrifuged 10mins at 15,000xg. Protein concentration as determined with Bradford reagent (BioRad) and an equal amount of input was used for control of RBM42 knockdown, typically approximately 2mg per sample. Cell extracts were then combined with the components for translation initiation: 1mM ATP, 10mM creatine phosphate, 1 mg/mL creatine phosphokinase, .02mM L-methionine, 12.5mM HEPES pH 7.4, 0.4mM cycloheximide, 0.25mM spermidine and 4mM GMP-PNP translation inhibitor. This reaction was incubated 17mins at 30C then loaded onto a 10-30% sucrose density gradient and run as described with the modification of 4hrs total centrifugation. The samples were then separated into 30 fractions to precisely isolate the unbound versus 48S RNA bound fractions. RNA was isolated from fractions 6-16, analyzed via qPCR comparing the relative mRNA in fractions 6-11 to 12-16.

### Immunofluorescence Staining of Patient Tissue

Slides with paraffin embedded sections were baked for 45mins in a 56C oven. The tissue was deparaffinized and hydrated with 2x xylene washes, 2× 100% ethanol, 2x 95% ethanol, 70% ethanol and 2x water. Then the samples underwent antigen unmasking with 10mins under pressure in citrate solution (10mM sodium citrate, 0.05% Tween 20, pH 6.0). The specimens were allowed to fully cool followed by 1x wash each in water and PBS. The samples were blocked in 10% goat serum and 0.3% Triton X-100 in PBS for 1hr at room temperature. Primary antibodies were diluted in diluent buffer (1% goat serum and 0.3% Triton X-100 in PBS) anti-RBM42 (Sigma #HPA043671) 1:100 and anti-cMyc (Abcam #ab32) 1:100 and incubated overnight at 4C. The next day the sections were washed 3x PBST (0.1% Tween-20). Secondary antibodies AF488 donkey anti-rabbit (Invitrogen) and CF568 goat anti-mouse (Biotium) diluted 1:300 in diluent buffer and incubated 1hr at room temperature. The samples were then washed 3x PBST and a final PBS wash. The slides were mounted with ProLong Gold antifade with DAPI. Images were acquired on the Zeiss LSM 900 confocal microscope at 20X magnification. Images were analyzed using ImageJ (Fiji). Fluorescence thresholds were set per patient and then low vs. high cells were counted manually.

### Immunohistochemistry (IHC)

The PDAC human tissue microarray was purchased from US Biolab (PAN040-01A). Tissue sections were deparaffinized and rehydrated through a series of ethanol washes as described for IF staining above. Following antigen retrieval in sodium citrate buffer, a standard immunohistochemistry protocol was done. Briefly, slides were incubated in methanol with 5% hydrogen peroxide, blocked with 2.5% normal goat serum (NGS) for 1 hour and incubated with primary antibody (RBM42, HPA043671, 1:400 dilution) overnight at 4°C. The following day, slides were washed with PBS, incubated with secondary antibody and stained with di-aminebenzidine (DAB) substrate kit (SK-4100, Vector Laboratories) for 10 minutes, washed, and then counterstained with hematoxylin (H-3401-500, Vector Laboratories). Finally, slides were dehydrated and mounted. Tissues were imaged using a KEYENCE BZ-X710 microscope. Brightfield images were captured in 10x and 20x magnification.

To quantify the RBM42 cytoplasmic and nuclear staining in the mouse tissues, ImageJ Fiji was used. First, to measure and subtract background signal a 10×10 square was drawn on an area of the image without staining. The same square was then drawn in the nucleus and cytoplasm separately and the mean intensity of RBM42 was measured. The background mean intensity was subtracted from the cytoplasm and nucleus mean intensities. N=3 brightfield images were taken per tissue per group (n=3 tissues, Groups = normal pancreas, PanIN, PDAC), and n=15-20 cells were quantified per image.

### Proximity Ligation Assay Immunofluorescence

Panc-1 cells seeded on 4-well Permanox chamber slides were fixed with 4% Paraformaldehyde at room temperature for 15 minutes, followed by quenching with 10mM Tris pH 7.4 and 3 washes with PBS. Blocking was done in 5% goat serum, 0.3% TritonX-100 in PBS for one hour at room temperature. Primary antibodies were diluted in diluent buffer (1% goat serum and 0.3% Triton X-100 in PBS) as follows anti-RBM42 (1:200; Sigma #HPA043671), anti-eIF2β (1:100; sc-166536 Santa Cruz Biotechnology), anti-RPS20 (1:500; 68270-1-Ig ProteinTech), anti-RPS3 (1:100; sc-376008 Santa Cruz Biotechnology) and incubated overnight at 4C. As a control, all antibodies were used alone (“none” conditions in figure) to quantify background signal. All samples were probed with anti-mouse MINUS and anti-rabbit PLUS Duolink secondaries. Duolink *In Situ* Red Reagents were used according to the manufacturers protocol (Millipore Sigma) and mounted with Duolink DAPI mounting media. Cells were imaged using a BC43 Spinning Disk Confocal microscope at 64x magnification. PLA particle/foci count was quantified using Fiji ImageJ and normalized to the number of nuclei per field.

### Tumorigenesis Studies

MiaPaca2 or BxPC3 (EV and Myc-HA) cells were transduced control or RBM42 depleting hairpins as described for the phenotyping assays above. Either 3×10^6^ cells for MiaPaca2 or 2×10^6^ cells for BxPC3 were suspended in a 1:1 solution of PBS and Matrigel (BD Biosciences) and injected into the subcutaneous space of athymic nude mice (Invigo). Once palpable, tumor volume was measure with electronic calipers and calculated using the formula (L/2)x(W/2)x(H.2)x(4/3)xπ. Animals were sacrificed before the max diameter of the tumor reached 30mm. Animal cohort numbers were chosen based on results from tumorigenesis pilot studies in mice. Researchers were not blinded to group identity.

### Quantification and Statistical Analysis

Graphed data are expressed as mean ± SEM unless indicated otherwise in the figure legends. The sample size (n) indicated in the figure legends represents biological replicates, experimental replicates or independent fields as appropriate and specified. Sample size was determined by the preliminary experimental results and no statistical methods were used to predetermine sample size. All samples meeting basic experimental conditions and baseline quality control were included in the graphed data and statistical analysis.

The statistical analyses described in each figure legend were performed using GraphPad Prism 10, significance was accepted at the 95% confidence level (∗ p < 0.05, ^∗∗^ p < 0.01, ∗∗∗ p < 0.001, ∗∗∗∗ p < 0.0001). The Xena browser^66^ was used PDAC normal versus tumor RNA expression data retrieval and statistical analysis. To compare quantitative protein expression between PDAC and normal pancreas, the UALCAN Proteomics online tool was used^88^. Otherwise, the statistical tests used are specified in the figure legend and are in agreement with the distribution of the data. The unpaired two-tailed t test was used when comparing two groups unless the variances were unequal when a Welch’s t test. For comparing multiple groups for polysome profiling qPCR, a two-way ANOVA with uncorrected Fisher’s LSD test was used. Alternatively, for comparing multiple groups ordinary 1-way ANOVA with Dunnett’s Multiple Comparison Test was used. This was dependent upon the interdependency of the groups or timepoints being compared. Gene ontology and other functional enrichment adjusted p values were determined using Fisher’s Exact Test with Bonferroni correction using Enrichr^89^.

## EXTENDED DATA FIGURE LEGENDS

**Extended Data Fig. 1:**
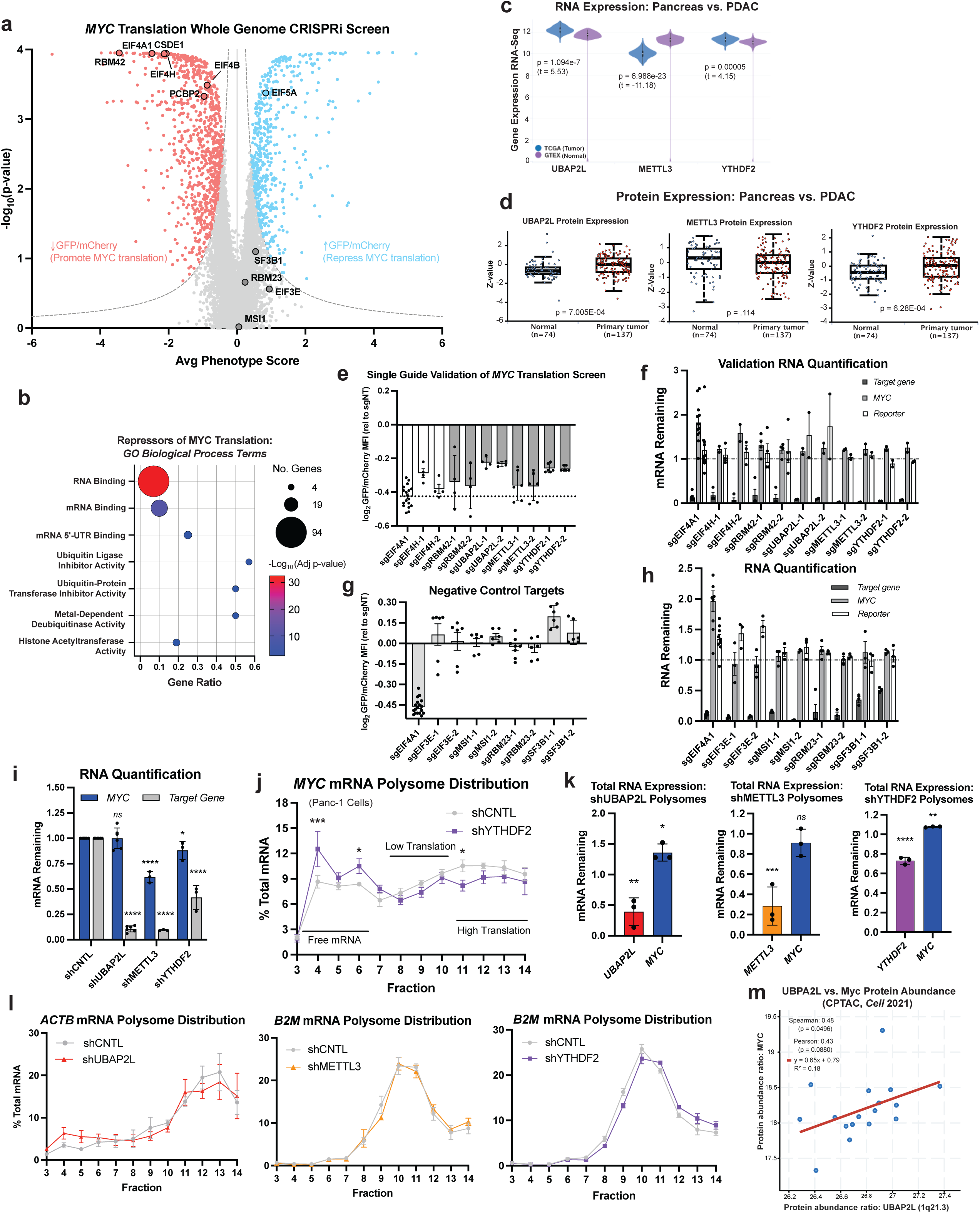
Validation and characterization of selective *MYC* translation screen candidates. **(a)** Volcano plot showing positive control known activators and negative control genes of *MYC* translation from Panc-1 CRISPRi reporter screen. eIF5A was previously identified and validated as the sole hit in a *MYC* translational repressor screen.^29^ **(b)** Gene ontology molecular function enrichment for the 309 candidate repressors of *MYC* translation in Panc-1 cells. Gene ratio is the fraction of genes identified within the GO term. **(c)** mRNA expression of *UBAP2L*, *METTL3* and *YTHDF2* in healthy pancreas (GTex) versus pancreatic adenocarcinoma (PDAC)^66^; Welch’s t-test. **(d)** Protein expression of UBAP2L, METTL3 and YTHDF2 normal versus tumor from CPTAC data set; two-sided t-test^45^. **(e)** Single guide validation with 2 independent sgRNAs in the Panc-1 *MYC* translational reporter cell line used for the CRISPRi screen at day 5 doxycycline treatment, positive controls *EIF4A1* and *EIF4H*, n ≥ 4. **(f)** qPCR RNA quantification of single guide validation experiments. **(g)** Single guide validation of negative control genes with 2 independent sgRNAs in the Panc-1 *MYC* translational reporter cells, n ≥ 6. **(h)** qPCR RNA quantification of negative control gene experiments. **(i)** Quantification of RNA from *UBAP2L*, *METTL3* and *YTHDF2* knockdown experiments in Panc-1 cells. **(j)** qPCR analysis of *MYC* mRNA from 10-50% sucrose gradient fractionation of control or YTHDF2 knockdown in Panc-1 cells, n=3. **(k)** Total RNA quantification from *UBAP2L*, *METTL3* and *YTHDF2* knockdown polysome experiments. **(l)** qPCR analysis of control genes, *ACTB* or *B2M*, from 10-50% sucrose gradient fractionation of control or *YTHDF2* knockdown in Panc-1 cells, n=3. **(m)** Correlation of UBAP2L and Myc protein abundance from human pancreatic adenocarcinoma specimens^45^, n=17. All graphs mean ± SEM. * p < 0.05, ** p < 0.01, *** p <0.001, **** p < 0.0001, *ns* = not significant (total RNA quantification 1-way ANOVA, uncorrected Fisher’s LSD test relative to own shCNTL & polysome distributions 2-way ANOVA, uncorrected Fisher’s LSD test). Extended data associated with Figure 1.

**Extended Data Fig. 2:**
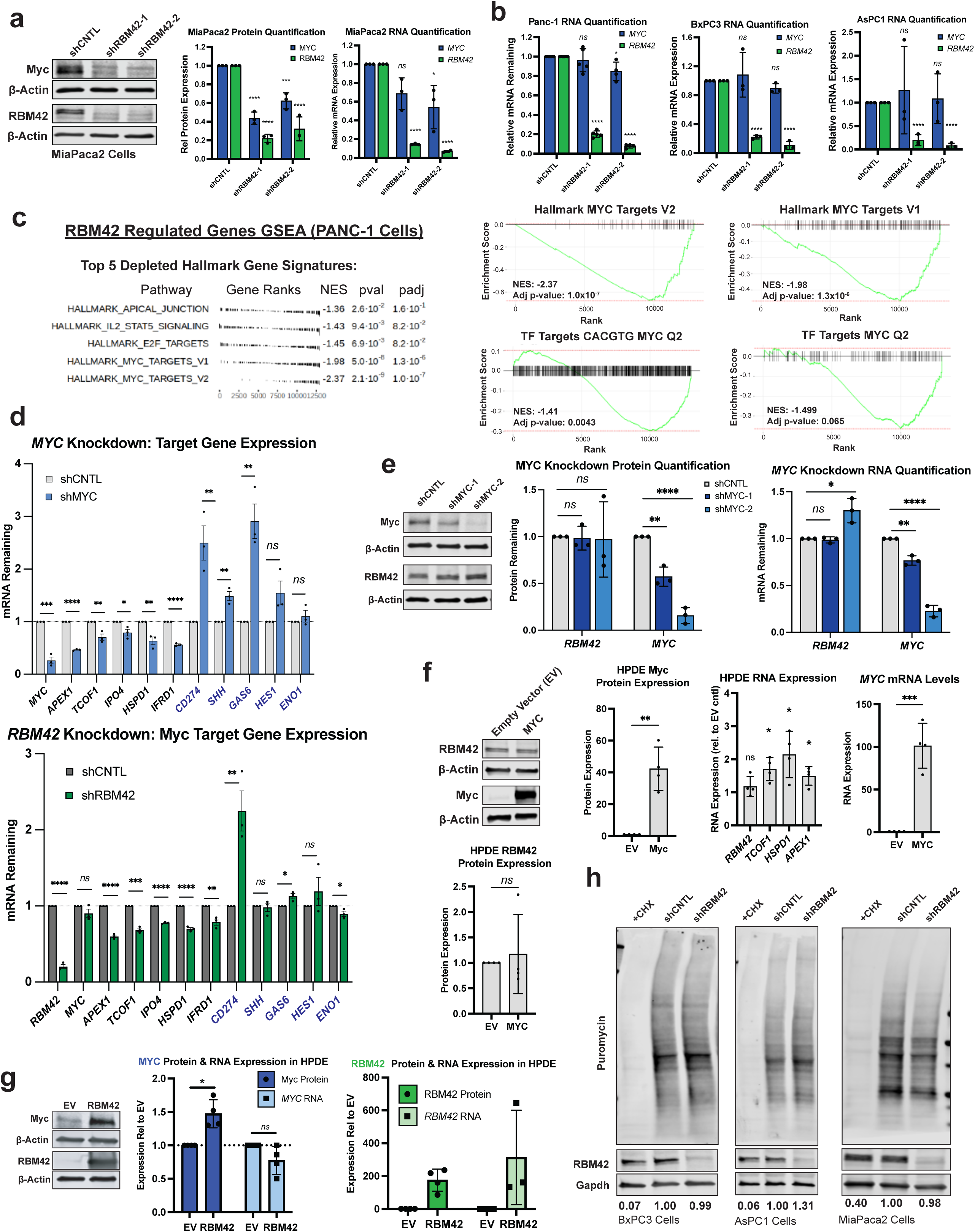
RBM42 is necessary and sufficient for regulating Myc protein expression. **(a)** Western blot analysis and quantification of RNA and protein expression of Myc and RBM42 with *RBM42* knockdown in MiaPaca2 pancreatic cancer cells, n=3. **(b)** qPCR RNA quantification for the knockdown experiments in Fig. 2d. PDAC cell line RNA and protein quantification 1-way ANOVA, uncorrected Fisher’s LSD test relative to own shCNTL. **(c)** Enriched Hallmarks and transcription factor (TF) targets from gene set enrichment analysis (GSEA) of genes with altered expression in Panc-1 shCNTL vs. shRBM42 RNA-seq data sets, Normalized Enrichment Score (NES). (**d**) qPCR quantification of RNA expression of core Myc targets and previously reported Myc targets in PDAC (blue text) in Panc-1 cells with *MYC* or *RBM42* depletion; mean ± SEM. Unpaired t-test relative to shCNTL. (**e**) Western blot analysis and quantification of RNA and protein expression of Myc and RBM42 with *MYC* knockdown in Panc-1 pancreatic cancer cells, n=3. Unpaired t-test. (**f**) Quantification and representative western blots of RBM42 and Myc RNA and protein levels with exogenous Myc expression in HPDE cells, n=4. Unpaired Welch’s t-test. **(g)** Quantification and representative western blots of RBM42 and Myc RNA and protein levels with exogenous RBM42 expression in HPDE cells, n=4. Unpaired Welch’s t-test. **(h)** Representative western blots of puromycin incorporation assay of global translation with RBM42 depletion in additional pancreatic cancer cell lines. Values indicate normalized puromycin incorporation relative to shCNTL. * p < 0.05, ** p < .01, *** p < 0.001, **** p < 0.0001, and *ns* = not significant. All graphs are mean ± SD unless otherwise indicated. Extended data associated with Figure 2.

**Extended Data Fig. 3:**
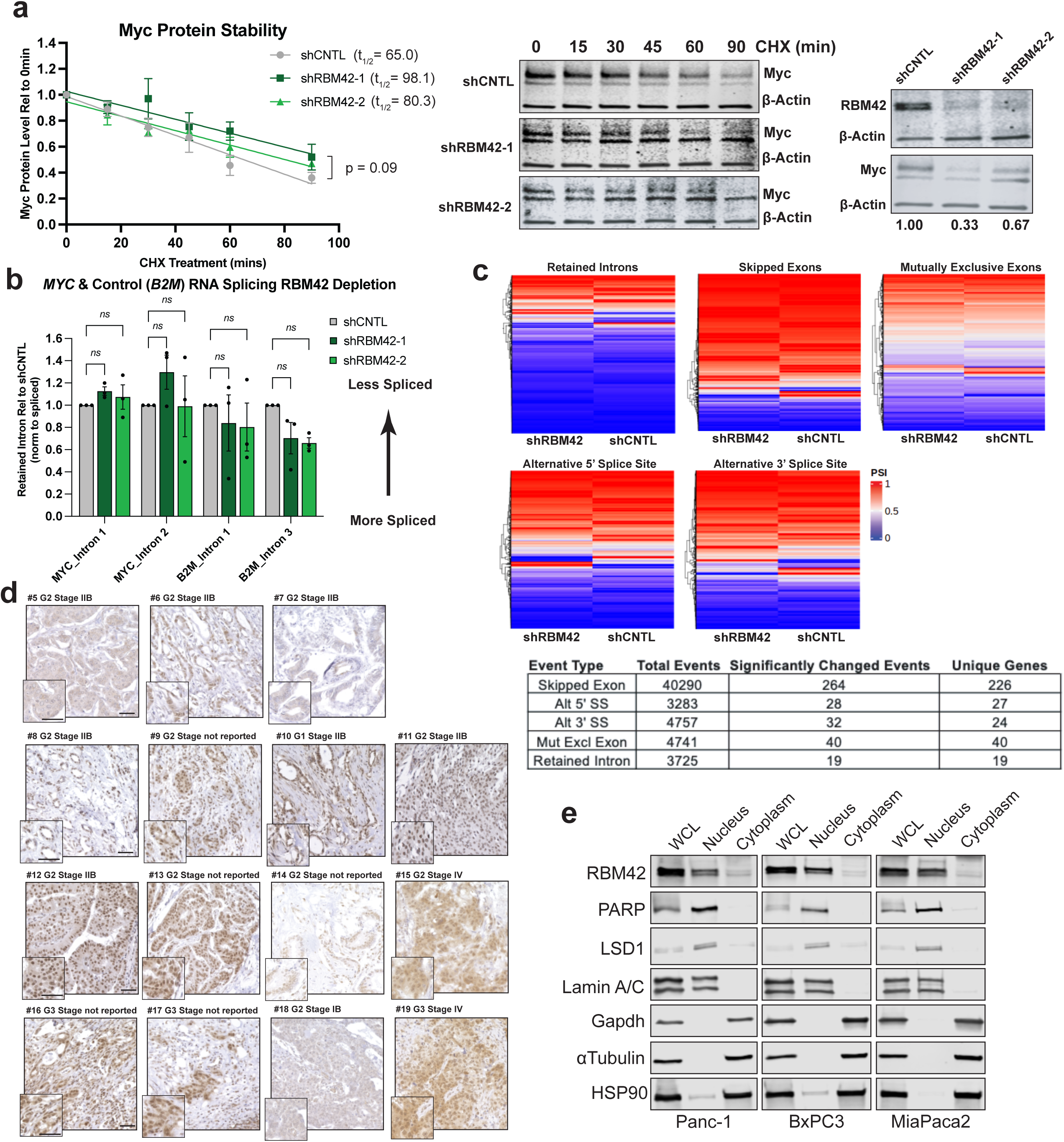
RBM42 loss does not alter Myc protein stability or RNA splicing. **(a)** Myc protein cycloheximide (CHX) stability assay quantification and representative western blots in Panc-1 cells, n=2. Both bands were quantified for Myc protein levels. P-value of the difference between the slopes. Values under blot indicate normalized Myc protein expression relative to shCNTL; mean ± SD. **(b)** qPCR quantification of retained introns for *MYC* and control mRNA *B2M*, n=3. 2-way ANOVA, uncorrected Fisher’s LSD test. **(c)** Panc-1 shCNTL and shRBM42 RNA-seq global splicing analysis (rMATS) collapsed to gene level, graphing the event with the highest ΔPSI, (n=3 replicates/condition). Table events FDR < 0.05 and |ΔPSI| > 0.1. (**d**) Additional human patient PDAC specimen RBM42 immunohistochemistry (IHC). Disease grade (G) and stage, where reported, are listed. Scale bar = 50μm. **(e)** Representative western blots of cytoplasmic and nuclear fractionation of pancreatic cancer cell lines. Graphs are mean ± SEM unless otherwise noted. ns = not significant. Extended data associated with Figure 2.

**Extended Data Fig. 4:**
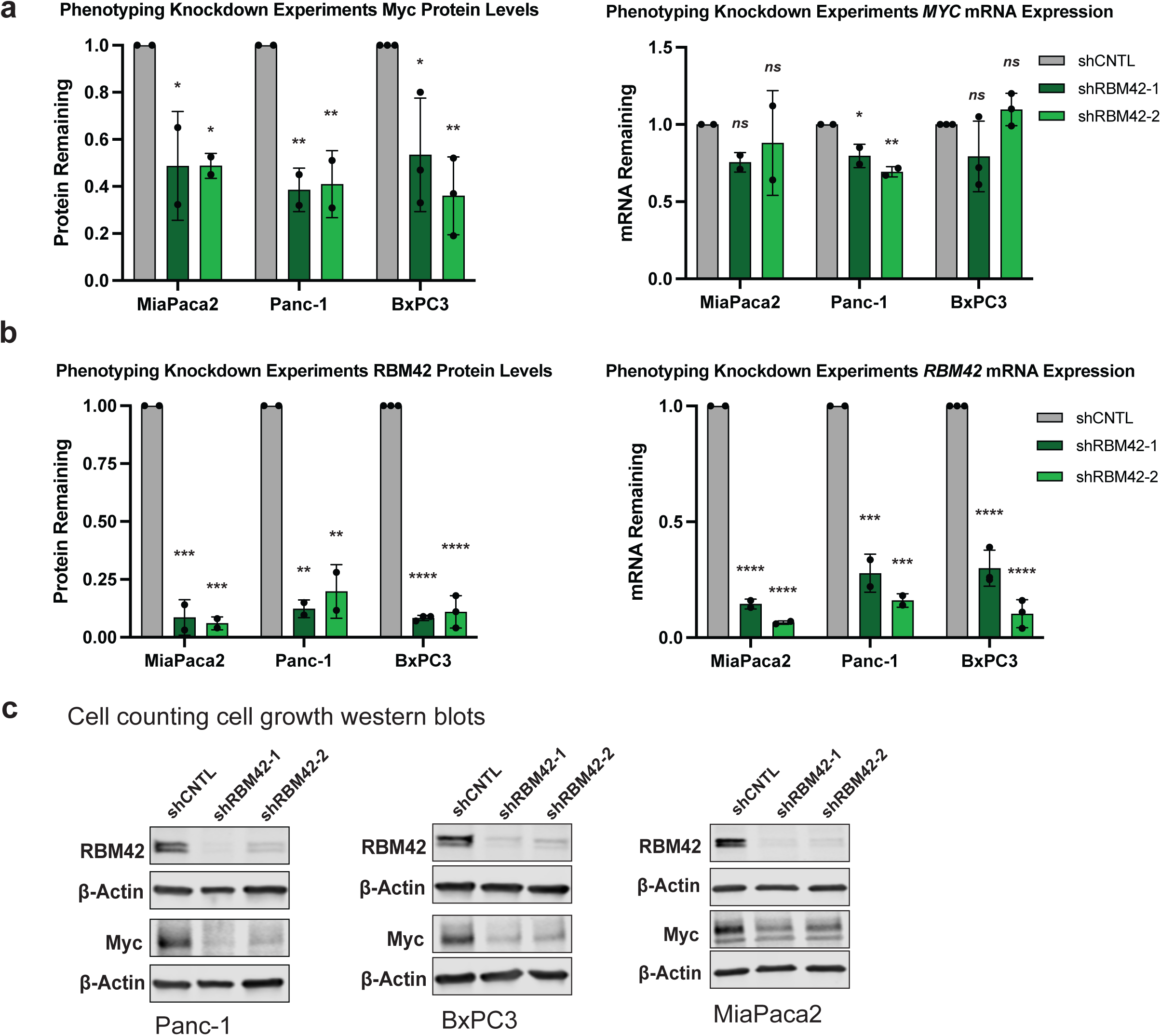
RBM42 and Myc RNA and protein expression in phenotyping experiments and validation of RBM42 necessity for PDAC cell growth. **(a)** Myc protein and RNA expression levels from the experiments in Fig. 3. **(b)** RBM42 protein and RNA expression levels from the experiments in Fig. 3. Graphs are mean ± SD; 1-way ANOVA, uncorrected Fisher’s LSD test relative to own shCNTL. **(c)** Representative western blots demonstrating for the PDAC cell line growth with RBM42 depletion. Graphs are mean ± SEM; unpaired t-test relative to shCNTL. * p < 0.05, ** p < 0.01, *** p < 0.001, **** p < 0.0001, *ns* = not significant. Extended data associated with Figure 3.

**Extended Data Fig. 5:**
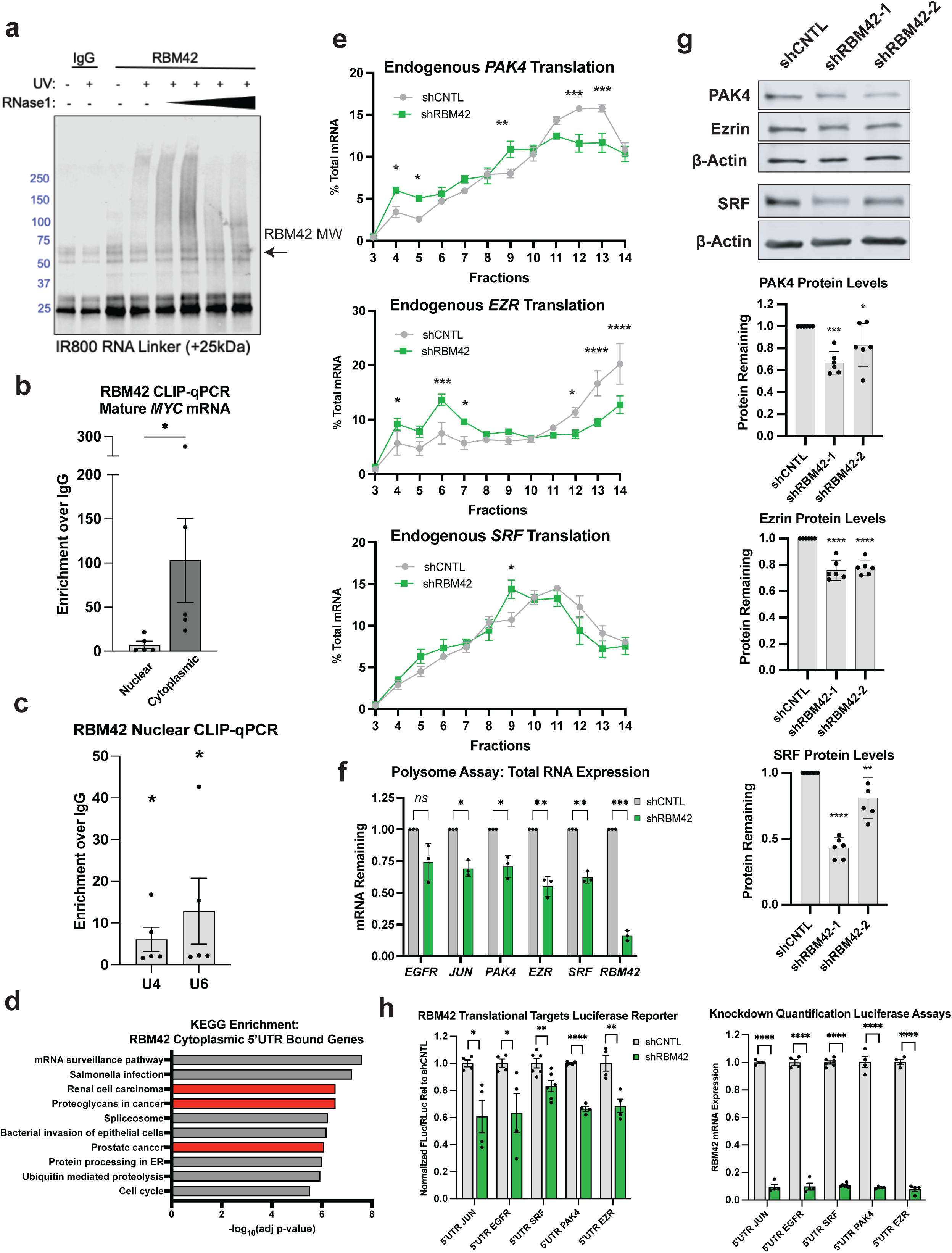
RBM42 regulates the translation of a broad suite of pro-oncogenic mRNAs. **(a)** Endogenous RBM42 irCLIP^73^ control RNA ligation blot -/+ UV crosslinking with titration of RNase I treatment. **(b)** Endogenous cytoplasmic or nuclear RBM42 CLIP-qPCR for mature *MYC* mRNA (lacking intron 2) fold over IgG control; n=5. Ratio paired t-test. **(c)** Nuclear RBM42 CLIP-qPCR positive control binding to known nuclear RNA interactions with U4 and U6 for the samples in (b); mean ± SEM. Ratio paired t-test compared to IgG control. **(d)** KEGG functional enrichment of genes bound in their 5’UTRs by RBM42. Terms related to cancer highlighted in red. **(e)** Polysome qPCR analysis of *PAK4*, *EZR* or *SRF* mRNA from 10-50% sucrose gradient fractionation of control or RBM42 knockdown in Panc-1 cells; n=3, mean ± SEM. 2-way ANOVA, uncorrected Fisher’s LSD test. **(f)** Total RNA quantification for the RBM42 knockdown polysomes; n=3. Paired t-test. **(g)** Representative western blots and protein quantification for PAK4, Ezrin (EZR), and SRF with RBM42 depletion in Panc-1 cells, n=6.1-way ANOVA, uncorrected Fisher’s LSD test. **(h)** Dual luciferase assays (DLA) with the 5’UTRs of additional RBM42 translational target genes with control or RBM42 depletion in Panc-1 cells. Renilla (RLuc) and firefly (FLuc) luciferase values normalized to RNA expression. RNA quantification validating *RBM42* depletion in the DLA samples; n=4-6, mean ± SEM. Unpaired t-tests. All graphs mean ± SD unless noted. * p < 0.05, ** p <0.01, *** p < .001, **** p < .0001, *ns* = not significant. Extended data associated with Figure 4.

**Extended Data Fig. 6:**
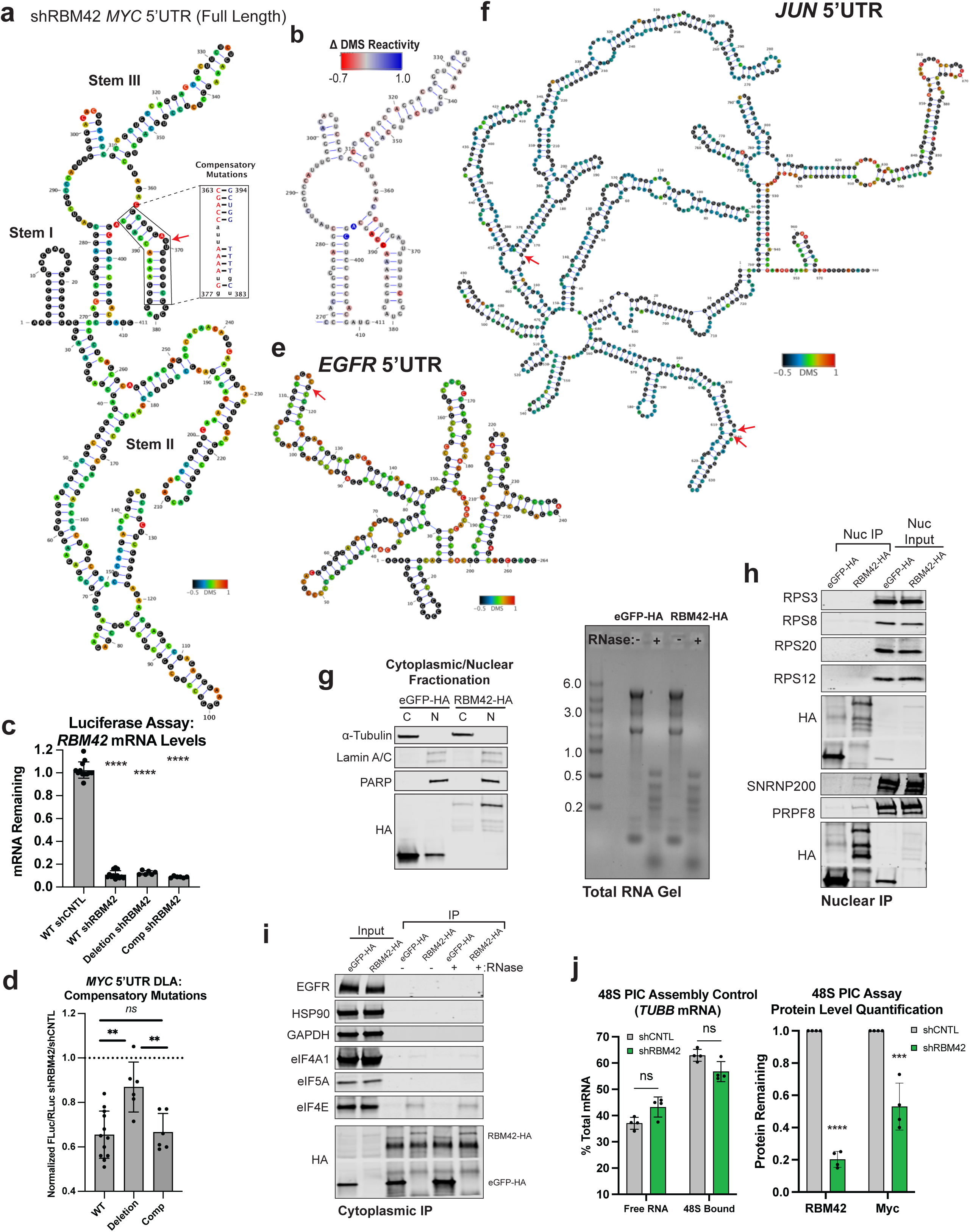
RBM42 alters *MYC* mRNA structure, interacts directly with 43S pre-initiation complex members in the cytoplasm and regulates *MYC* translation efficiency. **(a)** DMS-Seq determined full length *MYC* 5’UTR RNA structure in Panc-1 cells with RBM42 depletion. Red arrow indicates the RBM42 binding site. Box shows compensatory mutations red = 5’ mutations and blue = 3’ compensatory mutations to maintain structure. (**b**) *MYC* 5’UTR Stem III from shRBM42 structure showing the changes in DMS reactivity shRBM42 relative to shCNTL. **(c)** qPCR quantification of RBM42 mRNA levels in the dual luciferase assay experiments in Fig. 5c and ED Fig. 6d (1-way ANOVA, uncorrected Fisher’s LSD test to WT shCNTL). (**e**) DMS-Seq determined full length *EGFR* 5’UTR RNA structure in Panc-1 cells. Red arrow indicates the RBM42 binding site. (**f**) DMS-Seq determined full length *JUN* 5’UTR RNA structure in Panc-1 cells. Red arrows indicate the RBM42 binding sites. **(g)** Cytoplasmic and nuclear fraction blot to confirm proper fractionation for co-IP blots in Fig. 5f. Total RNA gel to confirm RNaseA digestion. **(h)** Representative IP-western blots of eGFP-HA or RBM42-HA from the nuclear fraction confirming known RBM42 interaction with SNRNP200 and PRPF8, but lack of interaction with 40S ribosomal components [corresponding fractionation blot in (g)]. **(i)** Representative IP-western blots of eGFP-HA or RBM42-HA from the cytoplasmic fraction probing for IP negative controls (EGFR, HSP90 and GAPDH) and initiation factors previously implicated in *MYC* translation initiation (eIF4A1, eIF5A and eIF4E). **(j)** 48S PIC assembly assay with control mRNA *TUBB* and quantification of decreased RBM42 and Myc protein levels for the experiment in Fig. 5h (Unpaired t-tests). ** p < 0.01, *** p < 0.001, **** p < 0.0001, ns = not significant. Extended data associated with Figure 5.

**Extended Data Fig. 7:**
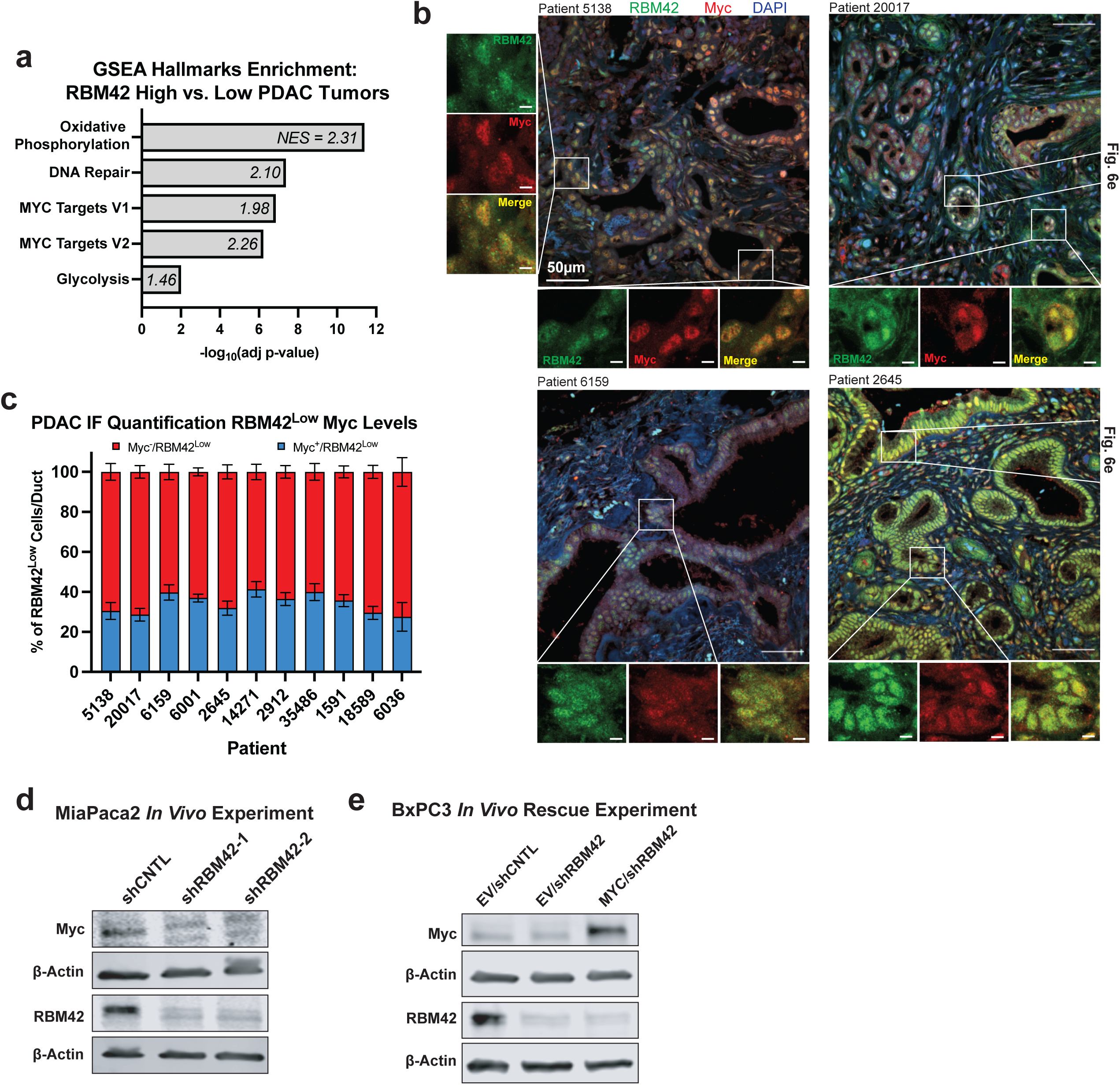
RBM42 regulates *in vivo* PDAC tumorigenesis through Myc. **(a)** Gene set enrichment analysis (GSEA) top 5 hallmarks comparing top quartile vs. bottom quartile of *RBM42* RNA expression in TGCA PAAD RNA-sequencing data (n=45 patients/quartile), Normalized Enrichment Score (NES). **(b)** Additional representative Myc and RBM42 immunofluorescence co-staining in human PDAC tissue. Location of higher magnification images shown in Fig. 6e indicated. Higer magnification images scale bar = 5μm. **(c)** Quantification of RBM42 low protein expression and Myc expression per cell overlap/patient; n = 6-48 ducts/patient, mean ± SEM. **(d)** MiaPaca2 and **(e)** BxPC3 Myc and RBM42 protein expression levels in the cells injected into subcutaneously into mice. Extended data associated with Figure 6.

